# Bmal1 integrates mitochondrial metabolism and macrophage activation

**DOI:** 10.1101/2019.12.20.884429

**Authors:** Ryan K. Alexander, Yae-Huei Liou, Nelson H. Knudsen, Kyle A. Starost, Chuanrui Xu, Alexander L. Hyde, Sihao Liu, David Jacobi, Nan-Shih Liao, Chih-Hao Lee

## Abstract

Metabolic pathways and inflammatory processes are under circadian regulation. While rhythmic immune cell recruitment is known to impact infection outcomes, whether the circadian clock modulates immunometabolism remains unclear. We find the molecular clock Bmal1 is induced by inflammatory stimulants, including Ifn-γ/lipopolysaccharide (M1) and tumor conditioned medium, to maintain mitochondrial metabolism under these metabolically stressed conditions in mouse macrophages. Upon M1 stimulation, myeloid-specific *Bmal1* knockout (M-BKO) renders macrophages unable to sustain mitochondrial function, enhancing succinate dehydrogenase (SDH)-mediated mitochondrial ROS production and Hif-1α-dependent metabolic reprogramming and inflammatory damage. In tumor-associated macrophages, the aberrant Hif-1α activation and metabolic dysregulation by M-BKO contribute to an immunosuppressive tumor microenvironment. Consequently, M-BKO increases melanoma tumor burden, while administrating an SDH inhibitor dimethyl malonate suppresses tumor growth. Therefore, Bmal1 functions as a metabolic checkpoint integrating macrophage mitochondrial metabolism, redox homeostasis and effector functions. This Bmal1-Hif-1α regulatory loop may provide therapeutic opportunities for inflammatory diseases and immunotherapy.

## INTRODUCTION

Inflammation and host defense are energetically costly processes that must balance the use of host resources with an efficient containment of infection or injury. This is underpinned by dynamic regulation of energy metabolism in immune cells in response to extrinsic signals, including cytokines, pathogen- and damage-associated molecular patterns and tumor-derived metabolites (Andrejeva & Rathmell, 2017; Buck, Sowell, Kaech, & Pearce, 2017; Ganeshan & Chawla, 2014; Hotamisligil, 2017; O’Neill, Kishton, & Rathmell, 2016). For instance, activation of macrophages by bacterial products, such as lipopolysaccharide (LPS) from gram-negative bacteria, shifts core metabolic function toward increased reliance on aerobic glycolysis with concomitant inhibition of mitochondrial respiration (Fukuzumi, Shinomiya, Shimizu, Ohishi, & Utsumi, 1996; Rodriguez-Prados et al., 2010; Tannahill et al., 2013). The depressed mitochondrial function appears to be by design, as this process serves multiple purposes. It leads to the so-called ‘broken TCA cycle’ due in part to shunting of citric acid to lipid synthesis (Andrejeva & Rathmell, 2017). Itaconate, also derived from citrate/aconitate, can modulate macrophage immune response through different mechanisms (Lampropoulou et al., 2016; Mills et al., 2018). By contrast, succinate accumulates through anaplerotic reactions, notably glutaminolysis (Tannahill et al., 2013). Succinate oxidation to fumarate mediated by succinate dehydrogenase (SDH)/ETC complex II activity is a primary source of mitochondrial reactive oxygen species (mROS) in inflammatory macrophages that are involved in bactericidal activity (Mills et al., 2016; West et al., 2011). Succinate/SDH is believed to trigger mROS production through accumulation of reduced coenzyme Q leading to reverse electron transfer to ETC complex I (Chouchani et al., 2014; Robb et al., 2018). These findings demonstrate a well-orchestrated metabolic signaling event at the expense of reduced fuel economy and compromised mitochondrial function in macrophages.

In addition to its bacterial-killing effect, mROS stabilize hypoxia-inducible factor (Hif) - 1α through inhibition of prolyl-hydroxylase enzymes that target Hif-1α for ubiquitination by the von Hippel–Lindau (Vhl) E3 ubiquitin ligase and subsequent proteasomal degradation (Bell et al., 2007; Jaakkola et al., 2001). Hif-1α is a master transcriptional regulator of genes involved in glycolysis and anabolic metabolism, thereby supplementing the energetic needs of the broken TCA cycle (Cramer et al., 2003; Masson & Ratcliffe, 2014; Semenza, Roth, Fang, & Wang, 1994). Hif-1α is also required for the expression of the urea cycle enzyme arginase-1 (*Arg1*). Arg1 and nitric oxide synthase 2 were initially designated as markers for M2 and M1 macrophages, as these two enzymes convert amino acid arginine to citrulline and nitric oxide, respectively. However, M1 activation also up-regulates *Arg1* through Hif-1α. Similarly, in the nutrient-deprived tumor microenvironment, tumor-derived lactate has been proposed to increase Hif-1α activity in tumor-associated macrophages (TAMs) to up-regulate *Arg1* (Colegio et al., 2014). Aberrant expression of Arg1 in TAMs results in local arginine depletion that inhibits antitumor immunity mediated by cytotoxic T cells and natural killer (NK) cells (Doedens et al., 2010; Steggerda et al., 2017). Accordingly, myeloid-specific deletion of *Hif1a* or *Arg1* suppresses tumor growth in mice (Colegio et al., 2014; Doedens et al., 2010). These observations suggest that the distinction between M1/M2 activation may not be as clear *in vivo* and highlight the importance of energetic regulation in immune cell activation.

The circadian rhythm has been implicated in many biological/pathological processes, including immune response and tumor progression (Hardin & Panda, 2013; Nguyen et al., 2013; Papagiannakopoulos et al., 2016). The molecular clock includes the master regulator Bmal1 (or Aryl hydrocarbon receptor nuclear translocator-like protein 1, Arntl) and its transcriptional partner Clock as well as the negative regulatory loop, including Nr1d1, Nr1d2, period (Per1/2/3) and cryptochrome (Cry1/2) proteins, and the positive regulator loop, including Rorα/β/γ (Hardin & Panda, 2013). Several nuclear receptors, such as peroxisome proliferator-activated receptors, Pparα, Pparδ/β and Pparγ, are downstream of Bmal1/Clock and control the expression of clock output genes (Canaple et al., 2006; S. Liu et al., 2013; Yang et al., 2006). The circadian clock is both robust and flexible. It has been demonstrated that time-restricted feeding in mice can synchronize the peripheral clock separable from the central clock (Damiola et al., 2000), suggesting that a primary function of circadian rhythm is to maximize the metabolic efficiency. In concert, we and others have shown that hepatic Bmal1 regulates rhythmic mitochondrial capacity in anticipation of nutrient availability (Jacobi et al., 2015; Peek et al., 2013). Prior studies have implicated the circadian oscillator in regulating macrophage inflammatory function. Notably, myeloid-specific *Bmal1* deletion disrupts diurnal monocyte trafficking and increases systemic inflammation and mortality in sepsis mouse models (Nguyen et al., 2013). Whether and how the circadian clock controls immune cell metabolism to modulate their effector functions remains unclear.

In the present study, we describe a cell-autonomous role for Bmal1 in macrophage energetic regulation. Bmal1 is induced following macrophage inflammatory stimulation. Its loss-of-function exacerbates mitochondrial dysfunction, energetic stress and Hif-1α-dependent metabolic reprogramming. By using the B16-F10 melanoma model, our results demonstrate that the regulatory axis between Bmal1 and Hif-1α dictates macrophage energy investment that is relevant for discrete activation/polarization states, including M1 and tumor-associated macrophages.

## RESULTS

### The circadian clock is a transcriptional module induced by M1 activation

To assess transcriptional regulators that modulate macrophage energetics and inflammatory function, we performed RNA sequencing (RNA-seq) comparing interferon-γ (Ifn-γ) primed bone marrow-derived macrophages without or with LPS stimulation (10 ng/mL for 8 hours, referred to as M1 activation). Gene ontology analysis using the DAVID platform was performed to identify clusters of transcription factors that were up- or down-regulated in inflammatory macrophages, which were used to generate a protein-protein interaction map using STRING (Figure 1 supplemental table 1 and Figure 1 supplemental figure 1A). Several activators of mitochondrial function/biogenesis were repressed, including *c-Myc* (Li et al., 2005), *Pparg*, Pparg co-activator 1 beta (*Pgc1b*), and mitochondrial transcription factor B1 (*Tfb1m*) and *Tfb2m*. On the other hand, the canonical inflammatory (e.g., *Nfkb1/2, Rela/b, Hif1a*, interferon regulatory factor 7 (*Irf7*) and *Irf8*) and stress response (e.g., *Atf3, Atf6b* and *Nfe2l2*) transcriptional modules were up-regulated. Interestingly, clusters of circadian oscillator components (e.g., *Per1, Cry1, Nr1d1, Nr1d2* and *Rora*) as well as nuclear receptors downstream of the molecular clock (e.g., *Ppard* and its heterodimeric partner *Rxra*) (S. Liu et al., 2013) were also induced.

We examined the expression of Bmal1, the non-redundant master regulator of circadian rhythm, and found that M1 activation (Figure 1A) or LPS treatment without Ifn-γ priming (Figure 1B) induced its mRNA and protein levels, peaking at 12 hours after the stimulation. Because LPS was directly added to the cell culture without changing the medium, the induction of Bmal1 was not due to serum shock (Tamaru et al., 2003). In fact, a one-hour LPS treatment in culture medium with 2% serum was sufficient to reset Bmal1 expression (Figure 1C), in a manner resembling serum shock (requiring a much higher serum concentration). Similar results were observed in mouse embryonic fibroblasts (MEFs), suggesting that the inflammatory regulation of Bmal1 was not macrophage-specific (Figure 1 supplemental figure 1B).

**Figure 1.**
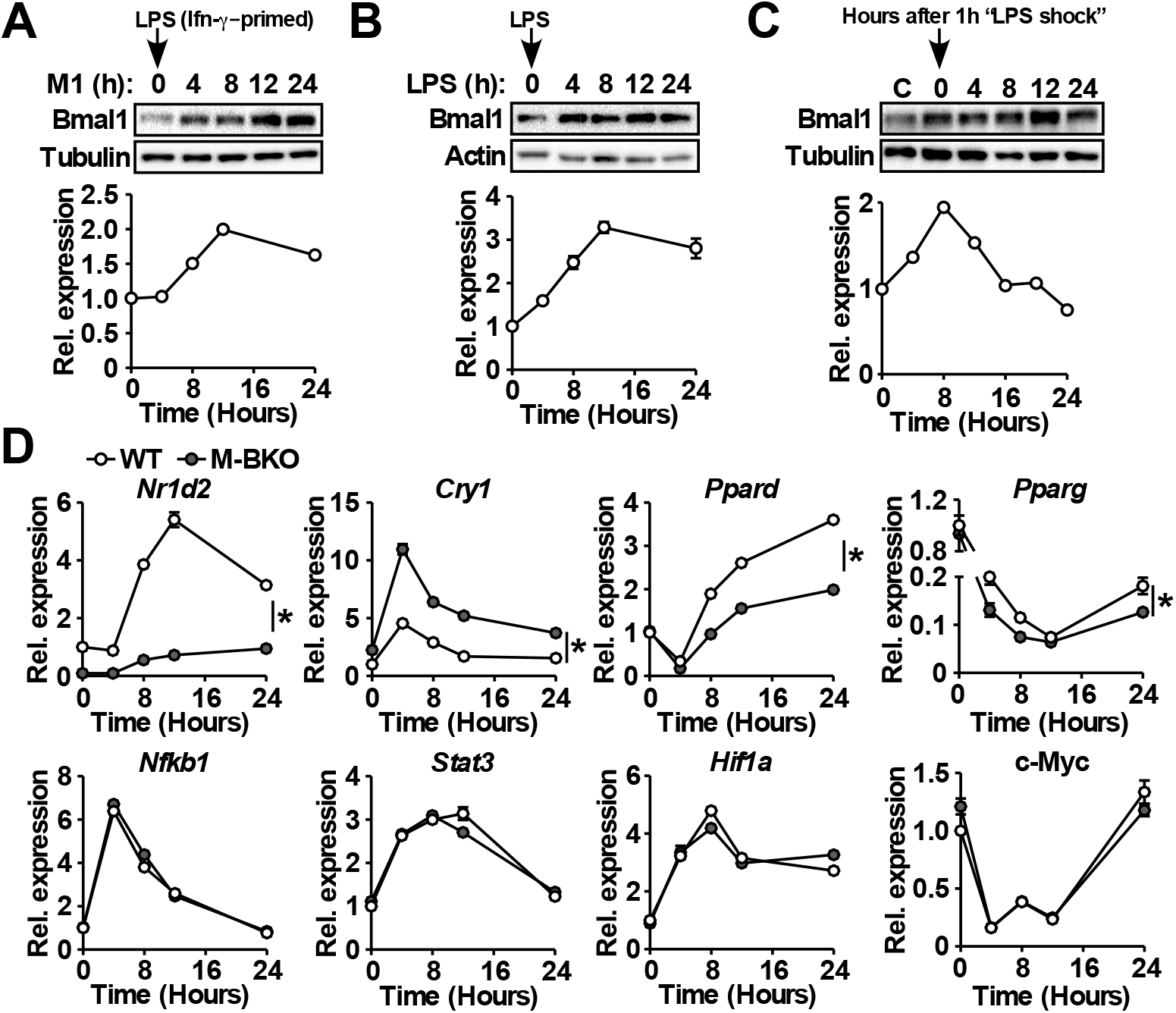
Macrophage Bmal1 is induced by M1 activation. (**A**), (**B**), and (**C**) Bmal1 protein levels (top) and relative gene expression determined by qPCR (bottom) in bone marrow-derived macrophages during a 24-hour time course of M1 activation (10 ng/ml Ifn-γ overnight priming + 10 ng/ml LPS) (A), treatment with LPS alone (100 ng/ml) (B), or acute LPS treatment for 1 hour (100 ng/mL) (C). For M1 and LPS only treatments, LPS was spiked in at time zero without medium change. For acute LPS treatment, cells were given with LPS for one hour followed by culture in DMEM, 2% FBS without LPS (time zero indicates medium change). N=3 biological replicates for qPCR. (**D**) Relative expression of circadian clock and inflammatory transcriptional regulators in M1-activated macrophages determined by qPCR. N=3 biological replicates, statistical analysis performed using 2-way ANOVA for WT vs. M-BKO across the time course. Data presented as mean ± S.E.M. *p<0.05. Experiments were repeated at least twice.

Myeloid-specific *Bmal1* knockout (M-BKO, *Bmal1f/f* crossed to *Lyz2-Cre; Bmal1f/f* was used as the wild-type control, WT) mice were generated to determine the role of circadian clock in macrophage function. M-BKO did not affect M1 induction of canonical inflammatory regulators, such as *Nfkb1, Stat3, Hif1a* and *c-Myc* (Figure 1D). The expression of genes downstream of Bmal1, including *Nr1d2, Cry1* and *Ppard*, was dysregulated, and there was a further reduction of *Pparg* expression by M1 activation in M-BKO macrophages compared to WT cells (Figure 1D). By contrast, M2 activation by Il-4 did not regulate *Bmal1* mRNA levels, and Il-4-induced expression of *Arg1* and *Mgl2* was not altered by M-BKO (Figure 1 supplemental figure 1C). These results suggest that the circadian clock may function as a downstream effector of M1 stimulation in a cell-autonomous manner.

### Bmal1 promotes mitochondrial metabolism in inflammatory macrophages

Because Pparδ/Pparγ are known regulators of mitochondrial function and energy substrate utilization in macrophages (Dai et al., 2017; Kang et al., 2008; Lee et al., 2006; Odegaard et al., 2007), we sought to determine the role of Bmal1 in macrophage bioenergetic control. In WT macrophages, M1 activation caused a progressive decrease in mitochondrial content, which was more pronounced in M-BKO macrophages (Figure 2A). The reduced mitochondrial content was likely due to mitophagy, as demonstrated by the increased level of the mitophagy receptor Bnip3 (Figure 2B). Measurement of ETC complex activity in isolated mitochondria indicated that M-BKO also caused a significant reduction in the activities of complex II and III, given an equal amount of mitochondrial protein, 6 hours after M1 stimulation (Figure 2C). Seahorse extracellular flux analysis showed that LPS injection increased the extracellular acidification rate (ECAR, indicative of lactic acid secretion) and decreased the oxygen consumption rate (OCR) of WT macrophages, as expected from aerobic glycolysis (Figure 2D). The ECAR and OCR were further enhanced and suppressed, respectively, in M-BKO macrophages. Similar results were obtained in thioglycollate-elicited peritoneal macrophages isolated from WT and M-BKO mice (Figure 2 supplemental figure 1A). By contrast, stable overexpression of *Bmal1* (Bmal1-OE) in RAW264.7 macrophages resulted in a higher OCR and lower ECAR after LPS stimulation, compared to control cells (Figure 2 supplemental figure 1B-C). To directly examine aerobic glycolysis, glucose was injected during the extracellular flux assay with or without co-injection with LPS. There was no difference in the basal glycolytic rate between WT and M-BKO macrophages (Figure 2E). LPS increased ECAR in both genotypes and to a greater extent in M-BKO macrophages. The induced ECAR could be blocked by injection of 2-deoxyglucose (2-DG), confirming the acidification was caused by aerobic glycolysis. Furthermore, an increase in the glycolytic rate was observed in splenic macrophages from M-BKO mice isolated 6 hours after *i.p*. injection of LPS, which was accompanied by lowered circulating glucose levels, indicative of increased glucose consumption by inflammatory myeloid cells in M-BKO mice, compared to WT animals (Figure 2 supplemental figure 1D-E).

**Figure 2.**
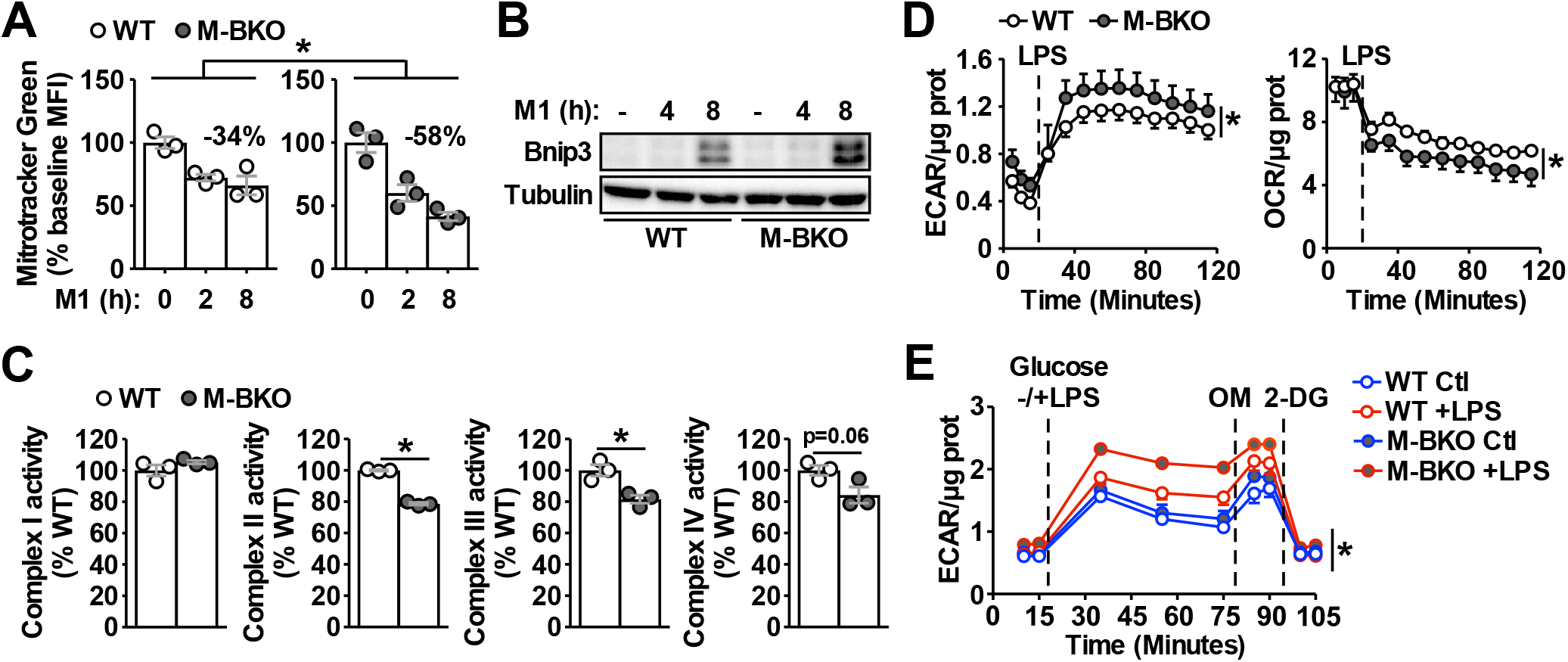
Bmal1 is required to maintain mitochondrial metabolism. (**A**) Assessment of mitochondrial mass in macrophages throughout a time course of M1 activation using Mitotracker Green (mean fluorescence intensity, MFI) determined by flow cytometry. N=3 biological replicates, statistical analysis performed using 2-way ANOVA for WT vs. M-BKO across the time course. (**B**) Immunoblot of the mitophagy regulator Bnip3 in M1-activated WT and M-BKO BMDMs. (**C**) Activities of ETC complexes in isolated mitochondria from WT and M-BKO macrophages after 6 hours M1 stimulation. N=3 biological replicates, statistical analysis performed using Student’s T test. (**D**) Extracellular flux analysis in Ifn-γ-primed macrophages measuring the changes in extracellular acidification rate (ECAR, left panel) and oxygen consumption rate (OCR, right panel) following LPS injection (100 ng/mL). Assay medium contained 5 mM glucose and 1 mM pyruvate in minimal DMEM with 2% dialyzed FBS, pH 7.4. N=5 biological replicates, statistical analysis performed using 2-way ANOVA for WT vs. M-BKO across the time course. (**E**) Glycolytic stress test in Ifn-γ-primed macrophages measuring ECAR following glucose (25 mM) injection, with or without LPS (100 ng/mL). Maximal glycolytic rate was determined by injection of oligomycin (OM, 2 μM), and glycolysis-dependent ECAR was determined by injection with 2-deoxyglucose (2-DG, 50 mM). Assay medium contained minimal DMEM with 2% dialyzed FBS, pH 7.4. N=5 biological replicates, statistical analysis performed using 2-way ANOVA for WT vs. M-BKO across the time course. Data presented as mean ± S.E.M. *p<0.05. Experiments were repeated at least twice.

To further assess the metabolic state, metabolomics analyses were employed to compare cellular metabolite levels of WT and M-BKO macrophages 0, 6 and 12 hours after M1 activation (Figure 3A-B and Figure 3 supplemental table 1). As has been reported (Tannahill et al., 2013), M1 activation caused accumulation of glycolytic intermediates (glucose-6-phosphate, fructose-6-phosphate and lactic acid) and depletion of TCA metabolites (e.g. citrate) but accumulation of succinate. Glycolytic metabolites and succinate were significantly higher in M-BKO macrophages compared to WT cells. M-BKO cells also showed accumulation of several amino acids and intermediates of the urea cycle (which detoxifies ammonia released from amino acid deamination) (Figure 3A and Figure 3 supplemental table 1). Consistent with the increased glycolytic metabolites, M1-stimulated glucose uptake and lactate production were higher in M-BKO macrophages (Figure 3C-D). These results suggest *Bmal1* loss-of-function leads to metabolic dysregulation in M1-stimulated macrophages.

**Figure 3.**
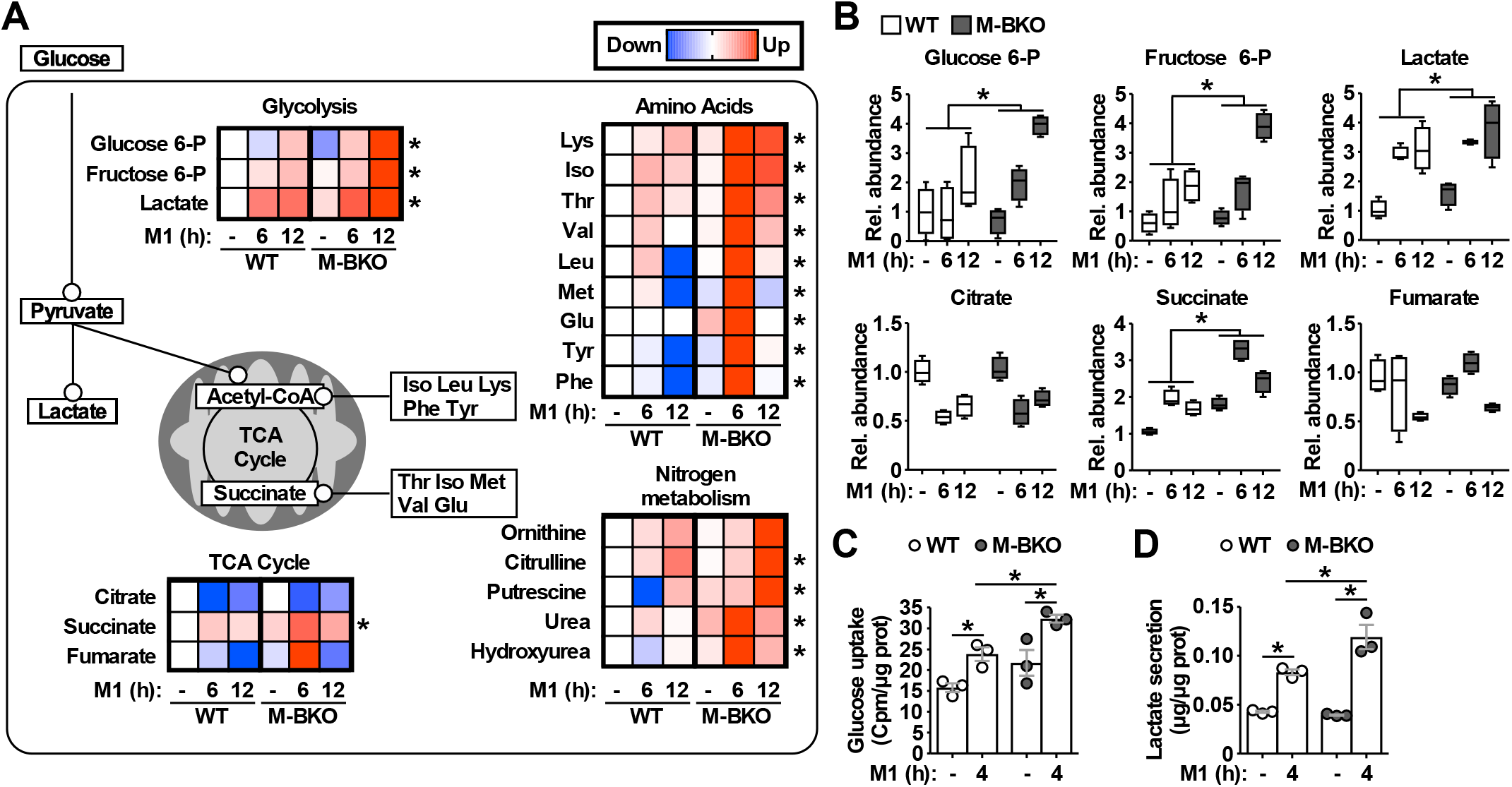
*Bmal1* deletion induces a metabolic shift for glycolytic and amino acid metabolism. (**A**) Summary of steady-state metabolomics data for differentially regulated metabolites from WT and M-BKO macrophages throughout a 12-hour M1 activation time course. Data presented as heat maps (normalized to WT control for each metabolite, each panel is the average of 4 biological replicates). Statistical analysis performed using 2-way ANOVA for WT vs. M-BKO across the time course. (**B**) Box plots of relative abundances for select metabolites in (A). (**C**) and (**D**) Uptake of [^3^H]-2-deoxyglucose and lactate secretion in control or M1-activated macrophages. N=3 biological replicates, statistical analysis performed using Student’s T test. Cell culture assays were repeated at least twice.

### The Bmal1-Hif-1α crosstalk regulates macrophage energy metabolism

As mentioned earlier, Hif-1α is a primary regulator of glucose metabolism in inflammatory macrophages. The enhanced aerobic glycolysis in M-BKO macrophages prompted us to examine whether Hif-1α activity was aberrantly elevated. Western blot analyses revealed that M1 activation led to a several-fold induction of Hif-1α protein levels in M-BKO macrophages compared to WT cells (Figure 4A), while Bmal1-OE RAW264.7 macrophages showed reduced Hif-1α protein (Figure 4 supplemental figure 1A). The expression of Hif-1α targets, such as lactate dehydrogenase A (*Ldha*), *Arg1* and *Il1b*, was enhanced by M-BKO and blocked by myeloid *Hif1a* knockout (M-HKO, Figure 4B). Hif-1α gene expression was not different between WT and M-BKO cells (Figure 1D). mROS derived from increased succinate oxidation has previously been demonstrated to stabilize Hif-1α protein in inflammatory macrophages (Mills et al., 2016). Metabolite analyses showed accumulation of succinate in M-BKO macrophages, suggesting that elevated mROS may be the cause of the increased Hif-1α protein. In fact, levels of mROS were higher in isolated mitochondria from M-BKO macrophages at 1 and 4 hours of M1 activation compared to WT macrophages (Figure 4C). Addition of succinate increased mROS production in mitochondria from both WT and M-BKO macrophages. An additional two-fold induction of mROS was detected in mitochondria from 4-hour M1 stimulated M-BKO, but not WT macrophages. Hif-1α protein accumulation could be normalized between genotypes by co-treatment with the antioxidant N-acetylcysteine (N-AC) or the competitive complex II inhibitor dimethylmalonate (DMM) that blocks mROS production (Figure 4D). Furthermore, M1-stimulated glucose uptake, lactate release and aerobic glycolysis were attenuated in myeloid-specific *Bmal1* and *Hif1a* double knockout macrophages (M-BHdKO, Figure 4 supplemental figure 1B-D), indicating the increased glucose utilization in M-BKO was Hif-1α-dependent.

**Figure 4.**
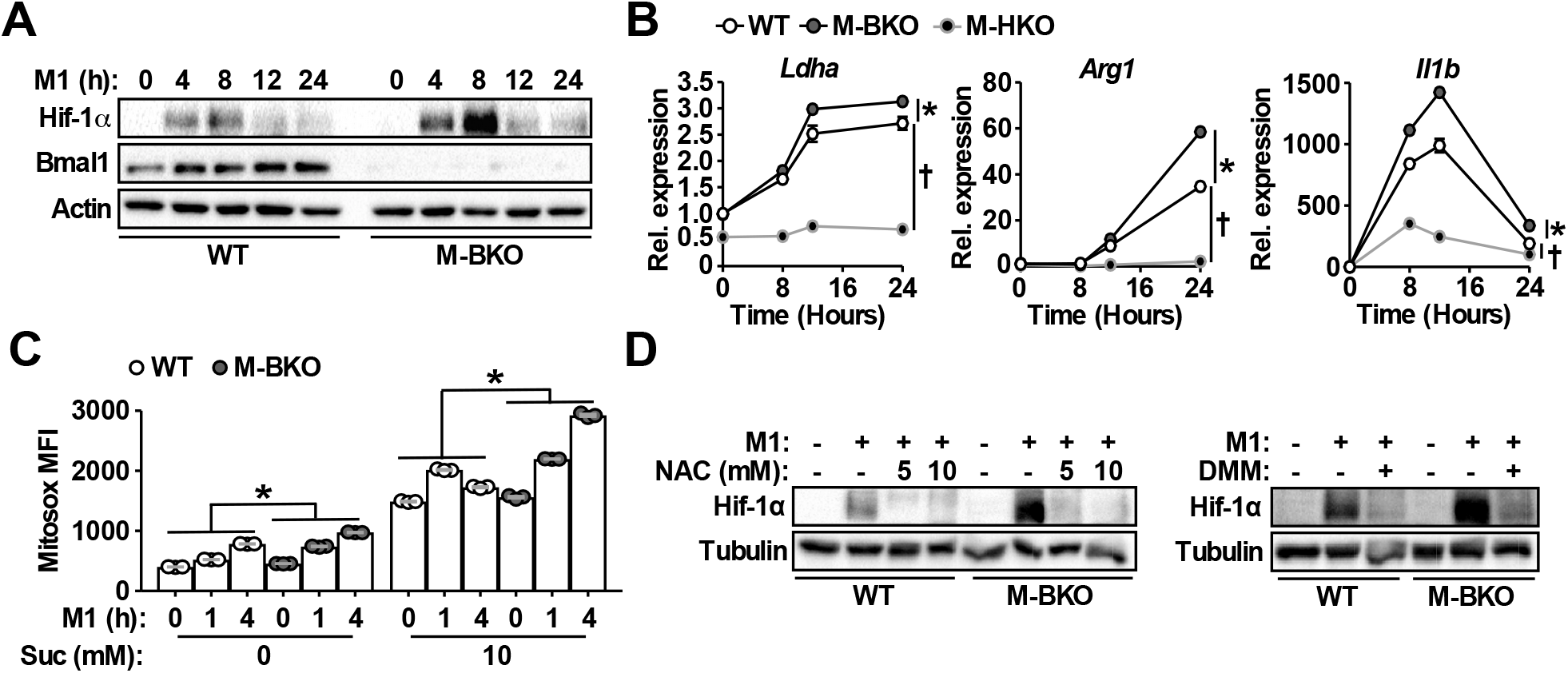
*Bmal1* loss-of-function increases oxidative stress and Hif-1α protein accumulation. (**A**) Immunoblots of Hif-1α and Bmal1 protein levels in WT and M-BKO macrophages during a 24-hour time course of M1 activation. (B) Relative expression of Hif-1α target genes in WT, M-BKO and M-HKO macrophages determined by qPCR. N=3 biological replicates, statistical analysis performed using 2-way ANOVA for WT vs. M-BKO or WT vs. M-HKO across the time course. (**C**) Measurement of mROS using MitoSox Red (mean fluorescence intensity, MFI) in mitochondria isolated from control or M1-activated macrophages. Succinate (Suc, 10 mM) was included during MitoSox Red staining where indicated. N=3 biological replicates, statistical analysis was performed using 2-way ANOVA for WT vs. M-BKO across the time course. (**D**) Hif-1α protein levels in control or 8 hours M1-activated macrophages co-treated with or without N-acetylcysteine (NAC) or 10 mM dimethylmalonate (DMM). Data presented as mean ± S.E.M. *p<0.05 for WT vs. M-BKO and **†** p<0.05 for WT vs. M-HKO. Experiments were repeated at least twice.

A previous study suggests that *Bmal1* deletion impairs the expression of *Nfe2l2* (which encodes Nrf2) and its downstream antioxidant genes thereby increasing oxidative stress (Early et al., 2018). However, we found that expression *Nfe2l2* and Nrf2-induced oxidative stress responsive genes, such as NAD(P)H quinone dehydrogenase 1 (*Nqo1*, Figure 4 supplemental figure 1E), were up-regulated in M-BKO macrophages upon M1 stimulation, suggesting that increased mROS associated with M-BKO was the cause rather than consequence of dysregulated Nrf2 signaling. Collectively, these data indicate that Bmal1 and Hif-1α regulate opposing metabolic programs and that Bmal1-mediated mitochondrial metabolism serves to fine-tune Hif-1α activity by modulating oxidative stress.

### *Bmal1* loss-of-function induces metabolic reprogramming toward amino acid catabolism

To fully characterize metabolic programs that were impacted by *Bmal1* loss-of-function, we compared RNA-seq data from control and M1-activated WT and M-BKO macrophages. These analyses revealed that the majority of M1-induced or suppressed genes were regulated in a similar manner between WT and M-BKO macrophages, suggesting that *Bmal1* gene deletion did not cause a general defect in inflammatory activation (Figure 5 supplemental figure 1A and Figure 5 supplemental table 1). Gene ontology analyses of the top enriched categories of M1-upregulated genes shared by both genotypes included regulation of apoptosis, response to stress and cytokine production. Among the top categories of suppressed genes were the cell cycle, DNA repair and carbohydrate metabolism. The latter showed that most TCA cycle enzymes were down-regulated by M1 activation (Figure 5A).

**Figure 5.**
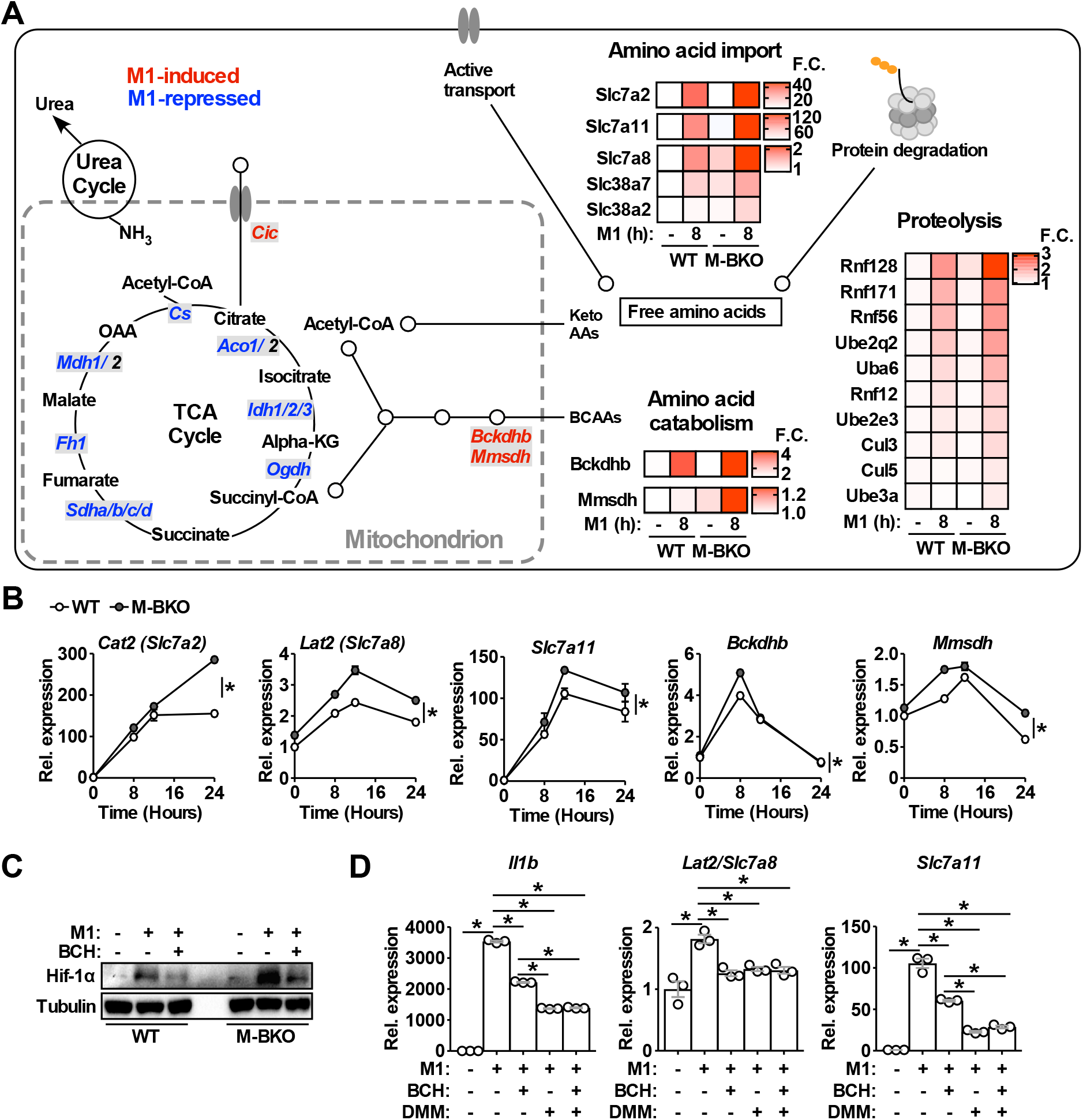
Genes involved in amino acid uptake and catabolism are up-regulated in M-BKO macrophages. (**A**) Schematic representation of M1-regulated genes involved in amino acid and TCA metabolism determined by RNA-seq. Genes in blue are downregulated while genes in red are upregulated by 8 hours M1 activation in both WT and M-BKO macrophages. Genes differentially regulated between genotypes are displayed in heat maps on the right. F.C: fold change. N=3 biological replicates. BCAAs: branch chain amino acids; Keto AAs: ketogenic amino acids; Cic: mitochondrial citrate carrier. (**B**) Relative expression of differentially regulated genes identified by RNA-seq and validated by qPCR in a 24-hour time course of M1 activation. N=3 biological replicates, statistical analysis performed using 2-way ANOVA for WT vs. M-BKO across the time course. (**C**) Hif-1α protein levels in control or 6 hours M1-activated macrophages with or without co-treatment of the neutral amino acid transport inhibitor 2-amino-2-norbornanecarboxylic acid (BCH, 10 mM). (**D**) Gene expression in control or 6 hours M1-activated macrophages with or without co-treatment of 10 mM BCH and/or the complex II inhibitor dimethyl malonate (DMM, 10 mM) determined by qPCR. N=3 biological replicates, statistical analysis performed using Student’s T test. Data presented as mean ± S.E.M. *p<0.05. Cell culture experiments were repeated at least twice.

Direct comparison between M1-stimulated WT and M-BKO macrophages revealed that genes more highly expressed in M1-activated M-BKO macrophages were enriched for protein catabolism and amino acid transport (Figure 5 supplemental figure 1B and Figure 5 supplemental table 1). These included genes encoding plasma membrane amino acid transporters (e.g., *Slc7a2, Slc7a8, Slc7a11, Slc38a2* and *Slc38a7*) as well as ubiquitin-activating, -conjugating and -ligating enzymes that target proteins for proteasomal degradation (e.g., ubiquitin-like modifier-activating enzyme 6 (*Uba6*), ubiquitin conjugating enzymes *Ube2q2* and *Ube2e3*, ring finger proteins *Rnf12, Rnf56, Rnf128*, and *Rnf171*, cullin 3 (*Cul3*) and *Cul5*, and ubiquitin protein ligase e3a (*Ube3a*)) (Figure 5A-B). The expression of enzymes involved in the breakdown of branched chain amino acids was also higher in M1-activated M-BKO cells, including branched chain keto acid dehydrogenase E1 subunit beta (*Bckdhb*) and methylmalonate semialdehyde dehydrogenase (*Mmsdh*). These results are consistent with increased amino acid catabolism observed in metabolite assays (Figure 3A). Interestingly, certain genes described above, notably *Slc7a8*, appeared to be counter-regulated by Hif-1α, as their induction by M1 stimulation was blunted in M-HKO macrophages (Figure 5 supplemental figure 1C).

Slc7a8, also called L-type amino acid transporter 2 (Lat2), transports neutral amino acids that could be converted to succinate and potentially contribute to Hif-1α protein stabilization. In line with increased amino acid metabolism, extracellular flux analysis showed that M-BKO macrophages showed enhanced glutamine utilization compared to WT cells, which was blocked by 2-amino-bicyclo-(2,2,1)-heptane-2-carboxylate (BCH), an L-type amino acid transporter inhibitor (Christensen, Handlogten, Lam, Tager, & Zand, 1969; Segawa et al., 1999) (Figure 5 supplemental figure 1D). BCH decreased and normalized levels of Hif-1α protein between WT and M-BKO macrophages (Figure 5C). In addition, treatment with either BCH or DMM suppressed the expression of *Il1b, Slc7a8* and *Slc7a11* induced by M1 stimulation (Figure 5D). The combination of BCH and DMM did not exert a greater effect over that of DMM alone. Thus, amino acid metabolism is up-regulated in response to dysregulated energy metabolism in M-BKO macrophages, which contributes to increased oxidative stress and Hif-1α activation.

### Macrophage *Bmal1* gene deletion promotes an immune-suppressive tumor-associated macrophage phenotype and enhances tumor growth

It has been suggested that myeloid-specific *Bmal1* deletion disrupts diurnal monocyte trafficking thereby increasing sepsis-induced systemic inflammation and mortality (Nguyen et al., 2013). Our results suggest that the cell-autonomous function of Bmal1 on macrophage metabolism and Hif-1α activation may contribute to the reported phenotype. Hif-1α regulates the polarization of M1 and tumor-associated macrophages, both of which are under energetically challenged conditions. We sought to determine whether the Bmal1-Hif-1α crosstalk plays a role in modulating TAM activation through a mechanism similar to that in M1 stimulation. Treatment of macrophages with conditioned medium from primary B16-F10 tumors (T-CM) increased the expression of *Bmal1* mRNA as well as protein (Figure 6A). M-BKO macrophages showed enhanced mROS production and Hif-1α protein induced by T-CM, compared to WT macrophages (Figure 6B-C). Tracking with Hif-1α stabilization, aerobic glycolysis was up-regulated by T-CM pretreatment in WT and to a greater extent in M-BKO macrophages (Figure 6D). T-CM elicited an energetic stress gene expression signature resembling M1 stimulation, including up-regulated amino acid metabolism (*Arg1*, *Slc7a8* and *Bckdhb*) and oxidative stress (*Slc7a11* and *Nqo1*) pathways in WT macrophages that were further induced by M-BKO (Figure 6E). Subsequently, we employed a mouse model of melanoma through subcutaneous injection of B16-F10 melanoma cells to assess the impact of myeloid *Bmal1* deletion on tumor growth. Tumor volume was increased in both male and female M-BKO mice compared to WT controls (Figure 6F). Furthermore, the expression of *Arg1, Slc7a8* and *Slc7a11* was up-regulated in F4/80^+^ cells isolated from tumors but not spleens of M-BKO mice compared to WT animals (Figure 6G). Of note, the mRNA levels of *Arg1, Slc7a8* and *Slc7a11* were substantially higher in tumor versus splenic F4/80^+^ cells, consistent with the results observed in T-CM treated macrophages.

**Figure 6.**
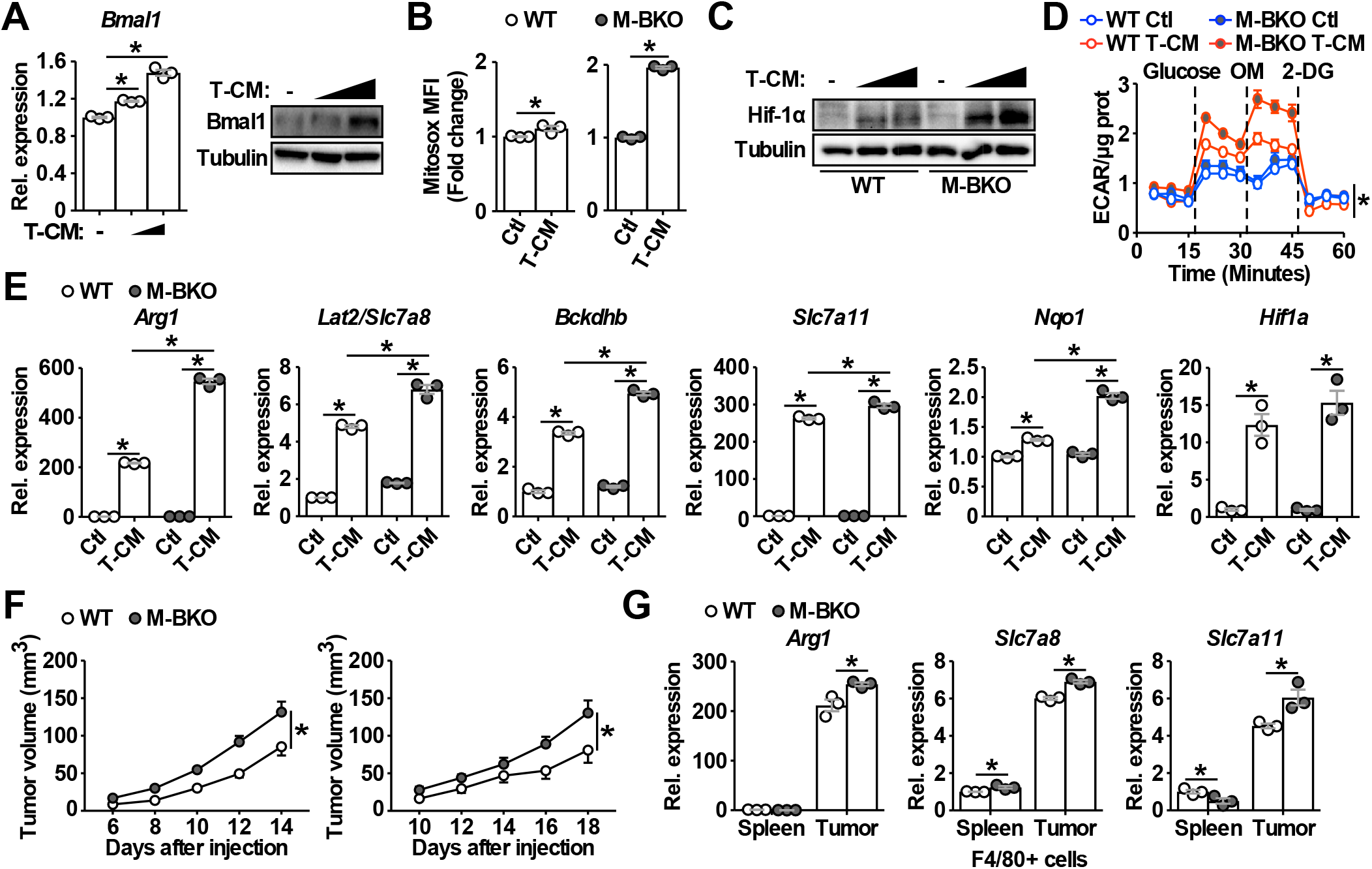
Bmal1 regulates tumor-associated macrophage polarization. (**A**) Bmal1 gene expression (left panel) and protein levels (right panel) in WT macrophages treated with control medium or increasing doses of B16-F10 tumor-conditioned medium (T-CM, diluted 1:3 or 1:1 with control medium) for 8 hours. N=3 biological replicates for qPCR, statistical analysis performed using Student’s T test. (**B**) Measurement of mROS using MitoSox Red (mean fluorescent intensity, MFI) in mitochondria from macrophages treated with control medium or T-CM diluted 1:1 with control medium for 1 hour. N=3 biological replicates, statistical analysis performed using Student’s T test. (**C**) Hif-1α protein levels in WT and M-BKO macrophages treated with control medium, T-CM diluted 1:1 with control medium, or undiluted T-CM for 4 hours. (**D**) Glycolytic stress test in macrophages pretreated with control medium or T-CM diluted 1:1 with control medium for 4 hours. . N=5 biological replicates. Statistical analysis performed using 2-way ANOVA comparing T-CM-treated M-BKO vs. WT cells across the time course. (**E**) Relative expression of genes involved in amino acid metabolism and oxidative stress response in macrophages treated with control medium or T-CM diluted 1:3 with control medium for 8 hours determined by qPCR. N=3 biological replicates, statistical analysis performed using Student’s T test. (**F**) Tumor volume in male (left) and female (right) WT and M-BKO mice. 300,000 B16-F10 cells were injected subcutaneously in the right flank. N= 18 (male) and 8 (female) mice, statistical analysis performed using 2-way ANOVA for WT vs. M-BKO mice across the time course. (**H**) Gene expression for F4/80^+^ cells isolated from B16-F10 tumors or spleens of female mice 14 days after injection. Tissues from 6 mice per genotype were pooled into 3 groups for leukocyte isolation. Statistical analysis performed using Student’s T test. Data presented as mean ± S.E.M. *p<0.05. Experiments were repeated at least twice.

To confirm that macrophage Bmal1 modulates tumor growth cell-autonomously and assess the effect of TAMs on anti-tumor immune response within the same host environment, we co-injected B16-F10 cells with either WT or M-BKO macrophages into the right or left flanks, respectively, of WT mice. Tumor growth rate was substantially higher when co-injected with M-BKO macrophages compared to WT cell co-injection (Figure 7A). In concert, co-injection with M-BKO macrophages led to a reduction in the CD8^+^ T cell population among tumor-infiltrating CD45^+^ leukocytes as well as functionally primed CD8^+^ T and NK cells that expressed Ifn-γ protein following stimulation with phorbol myristate acetate and ionomycin *ex vivo* (Figure 7B). Similar results were obtained when the co-injections were performed in M-BKO mice (Figure 7 supplemental figure 1A-B).

**Figure 7.**
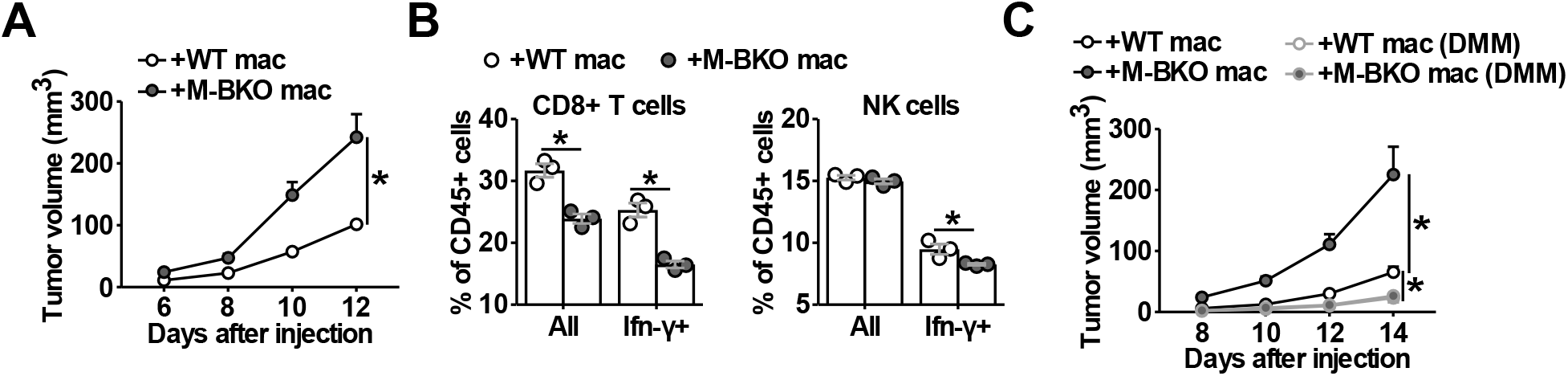
Macrophage Bmal1 modulates the antitumor activity. (**A**) Tumor volume in WT male mice co-injected with 500,000 B16-F10 cells and either 500,000 WT or M-BKO macrophages as indicated. N= 22 mice, statistical analysis performed using 2-way ANOVA for WT vs. M-BKO macrophage co-injection across the time course. (**B**) Flow cytometric analysis of tumorinfiltrating CD8^+^ T cells (CD45^+^CD3^+^CD8a^+^ cells, left panel) and NK cells (CD45^+^CD3^-^NK1.1^+^ cells, right panel) stimulated *ex vivo* with phorbol 12-myristate 13-acetate and ionomycin for Ifn-γ co-staining. Tumors from (A) were pooled into three groups prior to isolation of infiltrating leukocytes for flow cytometry. Statistical analysis performed using Student’s T test. (**C**) Tumor volume in WT male mice coinjected with 500,000 B16-F10 cells and either 500,000 WT or M-BKO macrophages supplemented without or with dimethylmalonate (DMM, approximately 150 mg/kg body weight per day in mouse diet). N=8 mice, statistical analysis performed using 2-way ANOVA for WT vs. M-BKO macrophage coinjection on control diet or for WT macrophage co-injection on the control diet vs. WT or M-BKO macrophage co-injection on the DMM-supplemented diet across the time course. Data presented as mean ± S.E.M. *p<0.05. Experiments were repeated at least twice.

We next sought to address the importance of oxidative stress in TAM activation. Similar to M1 macrophages, DMM blocked Hif-1α protein accumulation and attenuated *Arg1* up-regulation in T-CM-treated macrophages (Figure 7 supplemental figure 1C-D). Administering DMM (~150 mg/kg body) at the time of macrophage-tumor cell co-inoculation effectively suppressed melanoma tumor growth and normalized the difference in tumor promoting effects between WT and M-BKO macrophage (Figure 7C). These results reveal a unifying mechanism through which Bmal1 controls macrophage effector functions through bioenergetic regulation and suggest that targeting oxidative stress may provide a means to modulate the anti-tumor activity of TAMs.

## DISCUSSION

It has been reported that sepsis exerts a long-lasting effect on circadian rhythm alteration in mice (Marpegán, Bekinschtein, Costas, & Golombek, 2005; O’Callaghan, Anderson, Moynagh, & Coogan, 2012). In the current study, we show that inflammatory stimulants, including Ifn-γ/LPS and tumor-derived factors, control the expression of the circadian master regulator Bmal1 in the macrophages. Our data further demonstrate that Bmal1 is an integral part of the metabolic regulatory network and modulates macrophage activation, in part, through crosstalk with Hif-1α. The Bmal1-Hif-1α regulatory loop regulates the balance between oxidative and glycolytic metabolism in energetically stressed macrophages with distinct effector functions. *Bmal1* loss-of-function in M1-activated macrophages causes mitochondrial dysfunction, thereby potentiating mROS production and Hif-1α protein stabilization, which likely contributes to the increased sepsis-induced inflammatory damage reported for M-BKO mice (Nguyen et al., 2013). Within the tumor microenvironment, macrophage *Bmal1* gene deletion leads to compromised anti-tumor immunity and accelerated tumor growth in a mouse melanoma model. Therefore, the Bmal1-Hif-1α nexus serves as a metabolic switch that may be targeted to control macrophage effector functions.

Much attention has been focused on how inflammatory stimuli disrupt mitochondrial metabolism as a means to generate signaling molecules, including TCA metabolites and mROS. The analysis of transcriptional modules involved in macrophage inflammatory response reveals a coordinated effort in the control of mitochondrial activity. The expression of several regulators of mitochondrial biogenesis (e.g., *Pparg*) is down-regulated rapidly after M1 stimulation and rebounds between 8-12 hours, when *Bmal1* and *Ppard* expression are induced (Figure 1). Several lines of evidence indicate that Bmal1 plays a key role in restoring mitochondrial function and in modulating Hif-1α-mediated inflammatory response. The expression of transcription factors known to control mitochondrial bioenergetics discussed above (i.e., Pparγ and Pparδ) is down-regulated by M-BKO. Macrophages deficient in *Bmal1* are unable to sustain mitochondrial function upon M1 stimulation, while Bmal1 gain-of-function in RAW 246.7 macrophages promotes oxidative metabolism. The metabolic dysregulation in M-BKO macrophages further promotes Hif-1α-controlled glycolytic metabolism and other alternative sources for fuel utilization, such as amino acid catabolism. Although Bmal1 is best known for its role in circadian regulation and has been shown to control rhythmic monocyte recruitment and gene expression (Nguyen et al., 2013), our data suggest that LPS or M1 stimulation could “reset the clock” by inducing/resynchronizing the expression of *Bmal1*. In this context, Bmal1 controls the timing of glycolytic to oxidative metabolism transition that dictates the extent of Hif-1α activation and the associated inflammatory response.

Both Bmal1 and Hif-1α belong to the basic helix-loop-helix (bHLH) transcription factor family and have similar domain structures. However, they appear to regulate opposing metabolic programs, with Hif-1α serving as a master regulator of aerobic glycolysis and Bmal1 as a positive regulator of oxidative metabolism (Figure 7 supplemental figure 1E). The crosstalk between these two bHLH transcription factors is in part mediated by succinate and SDH/complex II-facilitated mROS production. Succinate is one of the entry points for anerplerosis that attempts to replenish TCA cycle metabolites depleted by disruption of mitochondrial oxidative metabolism. Increased protein/amino acid catabolism provides a source of anerplerotic reactions. Succinate accumulation and the subsequent oxidation to fumarate, however, generate mROS, which stabilizes Hif-1α protein to drive aerobic glycolysis. Our data suggest that amino acid metabolism appears to be down- and up-regulated by Bmal1 and Hif-1α, respectively, as demonstrated by the regulation of *Arg1* and *Slc7a8* gene expression. Hif-1α has also been shown to regulate Bnip3-mediated mitophagy that reduces mitochondrial oxidative capacity (Zhang et al., 2008). Therefore, Bmal1-controlled mitochondrial metabolism provides a break to this feedforward cycle to limit inflammatory damage. In line with this, previous work has demonstrated that myeloid *Bmal1* knockout mice have reduced survival rate upon *L. monocytogenes* infection (Nguyen et al., 2013). These observations indicate a tightly regulated metabolic program in the macrophage to execute effector functions and place Bmal1-regulated mitochondrial metabolism at the center of an orderly and balanced immune response.

Despite being characterized as M2-like, TAMs share several common features with M1-activated macrophages. Both of them function under nutrient-restricted conditions and Hif-1α is required for their activation. Previous studies implicate a glycolytic preference of TAMs in breast, thyroid, and pancreatic cancer (Arts et al., 2016; D. Liu et al., 2017; Penny et al., 2016). Our data confirm that T-CM treatment enhances glycolysis in the macrophage accompanied by increased Hif-1α protein (Figure 6C-D). *Arg1*, originally defined as an M2 marker, is a *bona fide* target of Hif-1α up-regulated in TAMs and M1 macrophages. Arg1 is involved in the urea cycle that detoxifies ammonia, and its induction supports amino acid catabolism. M-BKO macrophages show increased mROS, glycolytic metabolism and Hif-1α stabilization and up-regulation of *Arg1* and *Slc7a8* upon treatment with T-CM. Dysregulated amino acid metabolism has been shown to impact immune cell activation. Arginine depletion impairs lymphocyte function, as arginine is required for effector T cell and NK cell proliferation and maintenance (Geiger et al., 2016; Lamas et al., 2012; Steggerda et al., 2017). Slc7a8 transports neutral amino acids, including branched chain amino acids that are essential for lymphocyte activation and cytotoxic function (Sinclair et al., 2013; Tsukishiro, Shimizu, Higuchi, & Watanabe, 2000). As such, the increased amino acid utilization by M-BKO macrophages may contribute to the observed reduction in populations of Ifn-γ-producing CD8^+^ T and NK cells in tumor-infiltrating CD45^+^ leukocytes (Figure 7B and Figure 7 supplementary figure 1B). The fact that amino acid/protein metabolism and oxidative stress genes (*Arg1*, *Slc7a8* and *Slc7a11*) are up-regulated in TAMs, compared to splenic macrophages (Figure 6G) supports the notion that the energetic stress is also a key determinant of TAM polarization. As a proof-of-principle approach, we show that DMM treatment blocks T-CM induced Hif-1α protein stabilization *in vitro* and suppresses tumor growth *in vivo*. Thus, while M1 macrophages opt for an inefficient way to produce ATP, TAMs are limited in energy allocations. Both of these processes result in an energetically challenged state in which Bmal1-Hif-1α crosstalk controls the metabolic adaptation that shapes macrophage polarization. Future studies investigating mechanisms to harness this energetic stress will likely identify means to effectively modulate immune cell functions.

## MATERIALS and METHODS

### Reagents

Lipopolysaccharide, or LPS, from *Escherichia coli* strain K-235 (L2143) was from Sigma-Aldrich. Recombinant murine Ifn-γ (315-05) and Il-4 (214-14) were from Peprotech. The ETC complex II inhibitor dimethyl malonate (136441) and the L-type amino acid transport inhibitor 2-amino-2-norbornanecarboxylic acid, or BCH, (A7902) were from Sigma-Aldrich.

### Animals

All animal studies were approved by the Harvard Medical Area Standing Committee on Animal Research. Animals were housed in a pathogen-free barrier facility at the Harvard T.H. Chan School of Public Health. *Bmal1^fl/fl^* (stock # 007668), *Hif1a^fl/fl^* (stock # 007561), and *Lyz2-Cre* (stock # 004781) mice in the C57BL/6J background were obtained from Jackson lab and were originally contributed by Drs. Charles Weitz, Dmitriy Lukashev, and Irmgard Foerster, respectively. Floxed mice were crossed with *Lyz2-Cre* mice to generate myeloid-specific *Bmal1* and *Hif1a* knockout mice. Myeloid-specific *Bmal1* knockout mice were crossed with *Hif1a^fl/fl^* mice, and the resulting heterozygotes were crossed to generate myeloid-specific *Bmal1* and *Hif1a* double knockout mice. The genotypes were validated by both DNA genotyping and mRNA expression. Gender- and age-matched mice between 8-24 weeks of age were used for experiments. Similar results were obtained from male and female mice.

### Bone marrow-derived macrophage (BMDM) differentiation and cell culture

Macrophages were differentiated from primary mouse bone marrow from the femur and tibia using differentiation medium containing 30% L929-conditioned medium, 10% FBS, and pen-strep solution in low-glucose DMEM in 15 cm petri dishes. Media were changed every three days, and cells were lifted, counted, and plated in final format in tissue culture plates on days 7-8 of differentiation. For experiments, primary macrophages were maintained in low glucose DMEM containing 10% FBS and pen-strep. For M1 activation, macrophages were primed with 10 ng/mL Ifn-γ for 10-12 hours and subsequently stimulated with 10 ng/mL of *E. coli* LPS at the start of each experiment. Macrophages with Ifn-γ priming but without LPS were used as the control for M1 activation.

### Peritoneal and splenic macrophage isolation and culture

For peritoneal macrophage isolation, mice aged 2-4 months were *i.p*. injected with 3 mL of 3% thioglycollate (Sigma-Aldrich, T9032). After 3 days, mice were euthanized, and peritoneal cells were recovered by lavage. For isolation of splenic macrophages, mice were euthanized and spleens were dissected and mashed in growth medium (high glucose DMEM with 10% FBS) and passed through a 70 μm strainer. Cells were pelleted and resuspended in red blood cell lysis buffer. Monocytes and lymphocytes were recovered using the Ficoll-Paque Plus density gradient medium (GE Healthcare Life Sciences, 17144002) according to the manufacturer’s instructions, and suspension cells (lymphocytes) were washed away prior to experiments.

### LPS synchronization of Bmal1 expression

To synchronize Bmal1 gene and protein expression with LPS (or LPS shock), BMDMs or MEFs were given fresh culture medium with 2% FBS and 100 ng/mL LPS for 1 hour and then given fresh medium with 2% FBS without LPS. Bmal1 expression was tracked following LPS removal. For M1 or LPS induction of Bmal1 expression, cells were primed with or without 10 ng/mL Ifn-γ for 10-12 hours in DMEM, 10% FBS and subsequently stimulated with 10 ng/mL of LPS without changing the medium (time zero).

### Cell lines

Mouse embryonic fibroblasts (MEFs) were isolated from WT C57/BL6J mouse embryos and immortalized using the 3T3 protocol as previously described(Xu, 2005). For experiments, immortalized MEFs were maintained in growth medium containing high glucose DMEM, 10% FBS. RAW264.7 mouse macrophages (TIB-71) and B16-F10 mouse melanoma cells (CRL-6475) were purchased from ATCC. For generation of stable Bmal1 overexpressing RAW264.7 cells, the *Bmal1* coding sequence was cloned from mouse embryonic cDNA (forward primer: 5’ GGCGAATTCGCGGACCAGAGAATGGAC 3’; reverse primer: 5’ GGGCTCGAGCTACAGCGGCCATGGCAA 3’) and subcloned into the pBABE retroviral expression vector (Addgene, 1764). Retroviral vectors were transfected into Phoenix packaging cells, followed by collection of supernatants containing retroviruses. RAW264.7 macrophages were incubated with retroviral supernatants with 4 μg/mL polybrene, and infected cells were selected with 4 μg/mL puromycin. Control cells were transduced with the empty pBABE vector.

### Syngeneic tumor model and tumor measurement

Male and female WT and M-BKO mice aged 10-12 weeks were subcutaneously injected in the right flank with 300,000 B16-F10 mouse melanoma cells. For co-injection experiments, 500,000 B16-F10 cells were mixed with either 500,000 WT or M-BKO BMDMs (differentiation for 6 day) in the right and left flanks, respectively. Tumor dimensions were measured every two days by caliper after all mice had palpable tumors, and tumor volume was calculated as LxWxWx0.52 as previously described(Colegio et al., 2014). For DMM treatment, mice were switched to soft pellet, high fat diet (Bio-Serv, F3282) so that DMM can be mixed with the diet using a blender. The tumor growth rate was slower on high fat diet (Figure 7C) compared to normal chow (Figure 7A).

### RNA sequencing

RNA-seq was performed on RNA from 3 biological replicates per treatment. Sequencing and raw data processing were conducted at the Institute of Molecular Biology (IMB) Genomics Core and IMB Bioinformatics Service Core, respectively, at Academia Sinica (Taipei, Taiwan, ROC). In brief, RNA was quantified using the Quant-iT ribogreen RNA reagent (ThermoFisher, R11491), and RNA quality was determined using a Bioanalyzer 2100 (Agilent; RIN>8, OD 260/280 and OD 260/230>1.8). RNA libraries were prepared using the TruSeq Stranded mRNA Library Preparation Kit (Illumina, RS-122-2101). Sequencing was analyzed with an Illumina NextSeq 500 instrument. Raw data were analyzed using the CLC Genomics Workbench. Raw sequencing reads were trimmed by removing adapter sequences, low-quality sequences (Phred quality score of < 20) and sequences >25 bp in length and mapped to the mouse genome assembly (mm10) from University of California, Santa Cruz, using the following parameters: mismatches = 2, minimum fraction length = 0.9, minimum fraction similarity = 0.9, and maximum hits per read = 5. Gene expression was determined by the number of transcripts per kilobase million. Functional annotation clustering of differentially regulated genes was done using DAVID (https://david-d.ncifcrf.gov/), and the interaction maps of transcriptional regulators that were induced or repressed by M1 activation shown in Figure 1 supplemental figure 1A were generated using STRING (https://string-db.org/). Significantly changed genes were determined by p<0.05 and FDR<0.05.

### qPCR

Relative gene expression was determined by real-time qPCR with SYBR Green. The expression of the ribosomal subunit *36b4* (*Rplp0*) was used as an internal control to normalize expression data. Primer sequences are listed below:

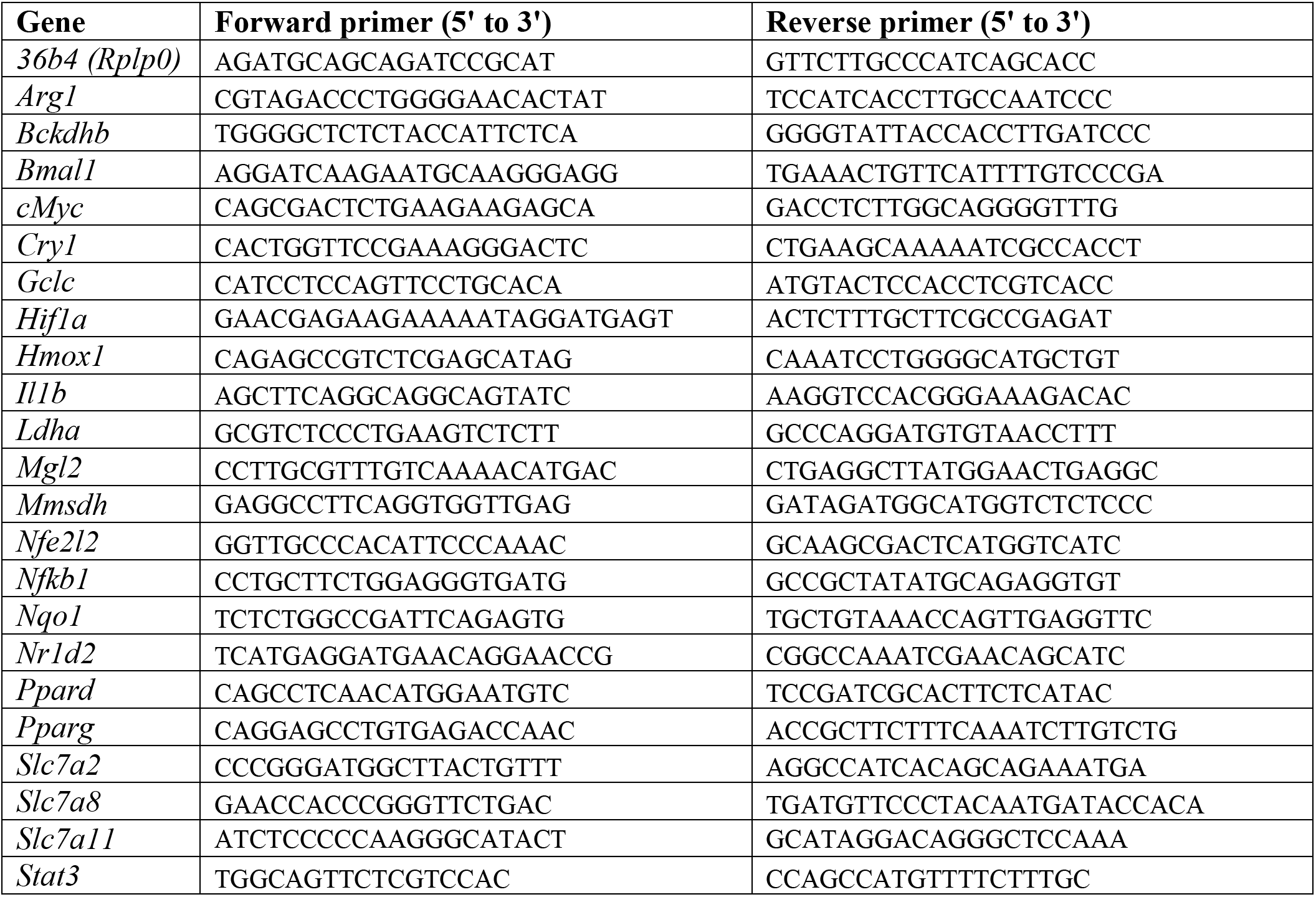

### Western blot

Standard Tris-Glycine SDS-PAGEs were run and transferred to PVDF membranes by wet transfer. Membranes were incubated with primary antibodies in TBST buffer with 1% BSA overnight. ECL signal was imaged using a BioRad ChemiDoc XRS+ imaging system. The antibody for Bmal1 (sc365645) was from Santa Cruz. The antibody for Hif-1α (NB100-449) was from Novus Biologicals. The antibodies for β-tubulin (2146) and β-actin (4970) were from Cell Signaling Technology.

### Extracellular flux analyses

Extracellular flux experiments were done using a Seahorse XF24 analyzer (Agilent) and FluxPaks (Agilent, 100850-001). 200,000 BMDMs, splenic/peritoneal macrophages or RAW264.7 cells were seeded into Seahorse XF24 plates for extracellular flux experiments. Minimal DMEM (pH 7.4) without phenol red and containing energy substrates as indicated was used as assay medium. 2% dialyzed FBS was added to media for experiments where LPS was injected during the assay to enhance responsiveness to LPS. Assay measurements were normalized to total protein content.

### Glucose uptake assay

BMDMs were plated at a density of 1 million cells per well in 12-well plates and stimulated as indicated. Cells were then washed with Krebs-Ringer bicarbonate HEPES (KRBH) buffer and then given 400 μL with KRBH buffer loaded with 0.8 μCi/well [^3^H]-2-deoxyglucose (PerkinElmer, NET549A001MC) and 0.5 mM unlabeled 2-deoxyglucose and incubated at 37°C for 30 minutes. 10 μL of 1.5 mM Cytochalasin B (Cayman Chemical, 11328) was then added to stop glucose uptake. 400 μL of lysate was used to measure levels [^3^H]-2-deoxyglucose by a scintillation counter, and the remaining lysate was used to measure total protein content for normalization.

### Measurement of lactic acid secretion

Lactic acid was measured in the supernatants of BMDMs using the Biovision Lactate Colorimetric Kit (K627) according to the manufacturer’s protocol. Readings were normalized to total cellular protein content.

### Mitochondrial isolation

Mitochondria were isolated from primary BMDMs by differential centrifugation. In brief, cells were resuspended in 500 μL of ice-cold mitochondrial isolation buffer consisting of 70 mM sucrose, 50 mM Tris, 50 mM KCl, 10 mM EDTA, and 0.2% fatty-acid free BSA (pH 7.2) and then extruded through 29-gauge syringes 20 times. Lysates were spun at 800g to pellet nuclei, and supernatants were spun at 8,000g to isolate mitochondria. Pelleted mitochondria were washed once more with 500 μL of mitochondria isolation buffer. Total mitochondrial protein content was determined by BCA assay.

### ETC activity assays in isolated mitochondria

The activities of ETC complexes I-IV were measured in isolated mitochondria using colorimetric assays as previously described (Spinazzi et al., 2012) with modifications. In brief, 15 μg of mitochondria were loaded per reaction for complexes III and IV, and 30 and 50 μg were used for complexes II and I, respectively. Complex I activity was determined by the decrease in absorbance at 340 nm corresponding to reduction of ubiquinone by electrons from NADH. Complex II activity was determined by the decrease in absorbance at 600 nm corresponding to reduction of decylubiquinone by electrons from succinate. Complex III activity was determined by the increase in absorbance at 550 nm corresponding to reduction of cytochrome C. Complex IV activity was determined by decrease in absorbance at 550 nm corresponding to oxidation of cytochrome C.

### Flow cytometry

For flow cytometry, BMDMs were seeded into low attachment plates for indicated treatments and resuspended by pipetting. Mitochondrial content in BMDMs was determined by flow cytometry of live cells stained with 100 μM Mitotracker Green FM (ThermoFisher, M7514) according to the manufacturer’s instructions.

For flow cytometry of tumor-infiltrating lymphocytes, cells were stimulated *ex vivo* with 20 ng/mL phorbol 12-myristate 13-acetate, or PMA, (Sigma-Aldrich, P8139) and 1 μg/mL ionomycin (Sigma-Aldrich, I0634) for 4 hours and co-treated with brefeldin A (Cell Signaling Technology, 9972) to inhibit cytokine release. Cells were stained with the fixable viability dye eFluor 455uv (ThermoFisher, 65-868-14) for 20 minutes at 4°C in FACS buffer (2% FBS and 1 mM EDTA in PBS), washed, and incubated with antibodies against indicated surface antigens for 30 min at 4°C. Cells were then washed twice and fixed with 2% paraformaldehyde for 1 hour at 4°C and resuspended and stored in FACS buffer prior to downstream analysis. Immediately before flow cytometric analysis, cells were permeabilized for intracellular staining using the Foxp3/Transcription factor staining buffer set (ThermoFisher, 00-5523-00) according to the manufacturer’s instructions. Of viable cells, CD8^+^ T cells were identified as CD45^+^ CD3^+^ CD8a^+^ cells and NK cells were identified by CD45^+^ CD3^-^ NK1.1^+^ staining. Antibodies for PerCp/Cy5.5-conjugated CD45 (103132), PE/Cy7-conjugated CD3e (100320), Alexa Fluor 700-conjugated CD8a (100730) and APC-conjugated NK1.1 (108710) were from Biolegend. The antibody for PE-conjugated Ifn-γ (12-7311-81) was from ThermoFisher Scientific.

To measure ROS production by isolated mitochondria, 15 μg of mitochondria were resuspended in 500 μL mitochondrial isolation buffer containing 5 μM MitoSox Red (ThermoFisher, M36008) and 100 μM MitoTracker Green FM with or without 10 mM sodium succinate. Mitochondria were incubated for 20 minutes at room temperature, washed with isolation buffer, and resuspended for flow cytometry. Mitochondria were identified by side scatter and positive MitoTracker Green staining for measurement of mean MitoSox Red intensity per population.

### Steady-state metabolomics

Untargeted metabolomics analysis using GC-TOF mass spectrometry was conducted by the West Coast Metabolomics Center at UC Davis. In brief, 10 million cells were lifted, pelleted, and washed twice with PBS for each replicate. Cell lysates were homogenized by metal bead beating, and metabolites were extracted using 80% methanol. Following extraction, cell pellets were solubilized using Tris-HCl Urea buffer (pH 8.0) containing 1% SDS to measure cellular protein content for each sample. All metabolite readings were normalized to total protein content.

### Collection of tumor-conditioned medium

Mice bearing subcutaneous B16-F10 tumors were sacrificed 20 days after injection with 500,000 cells. Tumors were dissected and weighed. Tumors were minced in growth medium containing 10% dialyzed FBS in high glucose DMEM (5 mL per gram of tissue) and incubated at 37°C for 2 hours. Conditioned medium was collected and filtered through 100 μm strainer followed by three spins at 1,000 rpm to pellet and remove residual cells/debris from the medium.

### Isolation of tumor-infiltrating immune cells

Subcutaneous mouse tumors were dissected, weighed, and then placed in 6-well plates with growth medium (RPMI, 5% FBS) and minced. Minced tissues were combined into three groups per genotype, spun down in 50 mL conical tubes, and resuspended in 20 mL digestion buffer (0.5 mg/mL collagenase IV, 0.1 mg/mL DNase I in HBSS medium). Tumors were digested at 37°C with gentle shaking for 30 minutes and vortexed every 10 minutes. Contents were filtered through a 100 mm mesh, and cells were pelleted and resuspended in 45% percoll in 1X HBSS and 1X PBS. Cells were spun at 2,000 rpm at 4°C with a swing bucket rotor for 20 minutes. The supernatant was aspirated, and the pellet was briefly resuspended in 5 mL ACK buffer to lyse red blood cells. Lastly, cells were pelleted and resuspended in growth medium for downstream applications.

To isolate F4/80^+^ cells from tumors, tissues were homogenized and processed as above to collect tumor-infiltrating leukocytes. F4/80^+^ cells were then isolated by positive selection using a rat anti-F4/80 antibody (Biolegend, 123120) and sheep anti-rat Dynabeads (ThermoFisher Scientific, 11035) according to the manufacturer’s instructions.

### Statistical analysis

All data are presented as mean ± SEM. GraphPad Prism 7 was used for statistical analyses. Two-tailed Student’s t test was used for comparisons of two parameters. Twoway ANOVA was used for multi-parameter analyses for time course comparisons. Cell-based experiments were performed with 3-5 biological replicates (cell culture replicates). For tumor volume, outliers were determined using a Rout test (p<0.05), and outliers were omitted from downstream experiments.

## ACKNOWLEDGEMENTS

We thank Drs. D. Cohen, D. Sinclair, and P. Weller for critical comments, A. Yesian, L. Dai, P. Basak, R. Chen, P. Hu, and F. Onal for technical help and the Academia Sinica (Taipei, Taiwan, ROC) for RNA sequencing/data analysis (IMB Genomics Core and Bioinformatics Service Core) and grant support (AS-106-TP-L08 to N.S.L: AS-106-TP-L08-1 and AS-106-TP-L08-3). Y.H.L. was supported by funds from Ministry of Science and Technology, Taiwan, ROC. This work was supported by grants from NIH (F31GM117854 to R.K.A.; F31DK107256 to N.H.K.; R01DK113791 and R21AI131659 to C.H.L) and American Heart Association (16GRNT31460005 to C.H.L.).

## AUTHOR CONTRIBUTIONS

R.K.A., Y.H.L., N.H.K., K.A.S., and C.X. performed the experiments. A.L.H., S.L., and D.J. generated reagents. N.S.L. assisted with RNA-seq data analysis. R.K.A. and C.H.L. conceptualized the study, designed experiments, interpreted data, and wrote the manuscript. C.H.L. supervised the study.

## COMPETING INTERESTS

The authors declare no competing interests.

---

**Figure 1 supplemental figure 1.**
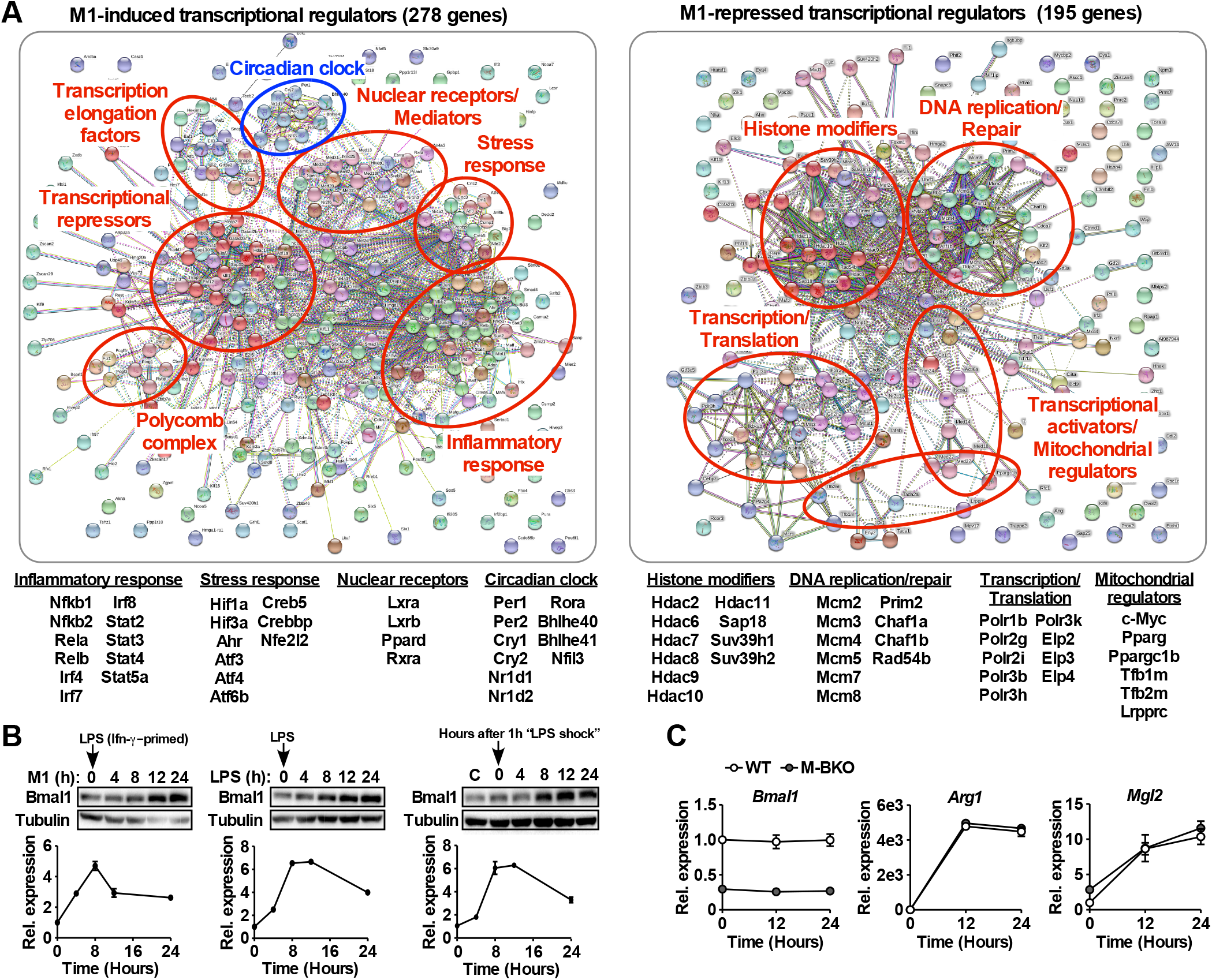
The circadian clock is a transcriptional module induced by M1 activation. (**A**) Annotated protein-protein interaction maps generated by STRING of transcriptional regulators that were significantly induced (left) or repressed (right) in WT macrophages by 8 hours M1 activation (Ifn-γ priming followed by stimulation with 10 ng/mL LPS) compared to time-matched controls (Ifn-γ priming without LPS), determined by RNA-seq (p<0.05, FDR<0.05, |F.C.|>1.5). N=3 biological replicates. F.C.: fold change. See Supplemental table 1 for the complete gene list. (**B**) Bmal1 protein levels (top panels) and relative gene expression determined by qPCR (bottom panels) in mouse embryonic fibroblasts during a 24 hour time course of M1 activation (10 ng/ml Ifn-γ overnight priming + 10 ng/ml LPS, left panels), treatment with LPS only (100 ng/ml, middle panels), or 1-hour acute LPS treatment (100 ng/mL, right panels). For M1 and LPS only treatment, LPS was spiked in at time zero without medium change. For acute LPS treatment, cells were given with LPS for one hour followed by culture in DMEM, 2% FBS without LPS (time zero indicates medium change). N=3 biological replicates for qPCR. (**C**) Gene expression during a 24-hour time course of Il-4 treatment in WT and M-BKO macrophages determined by qPCR. N=3 biological replicates. Data presented as mean ± S.E.M. *p<0.05. Cell culture experiments were repeated at least twice.

**Figure 1 supplemental table 1.**
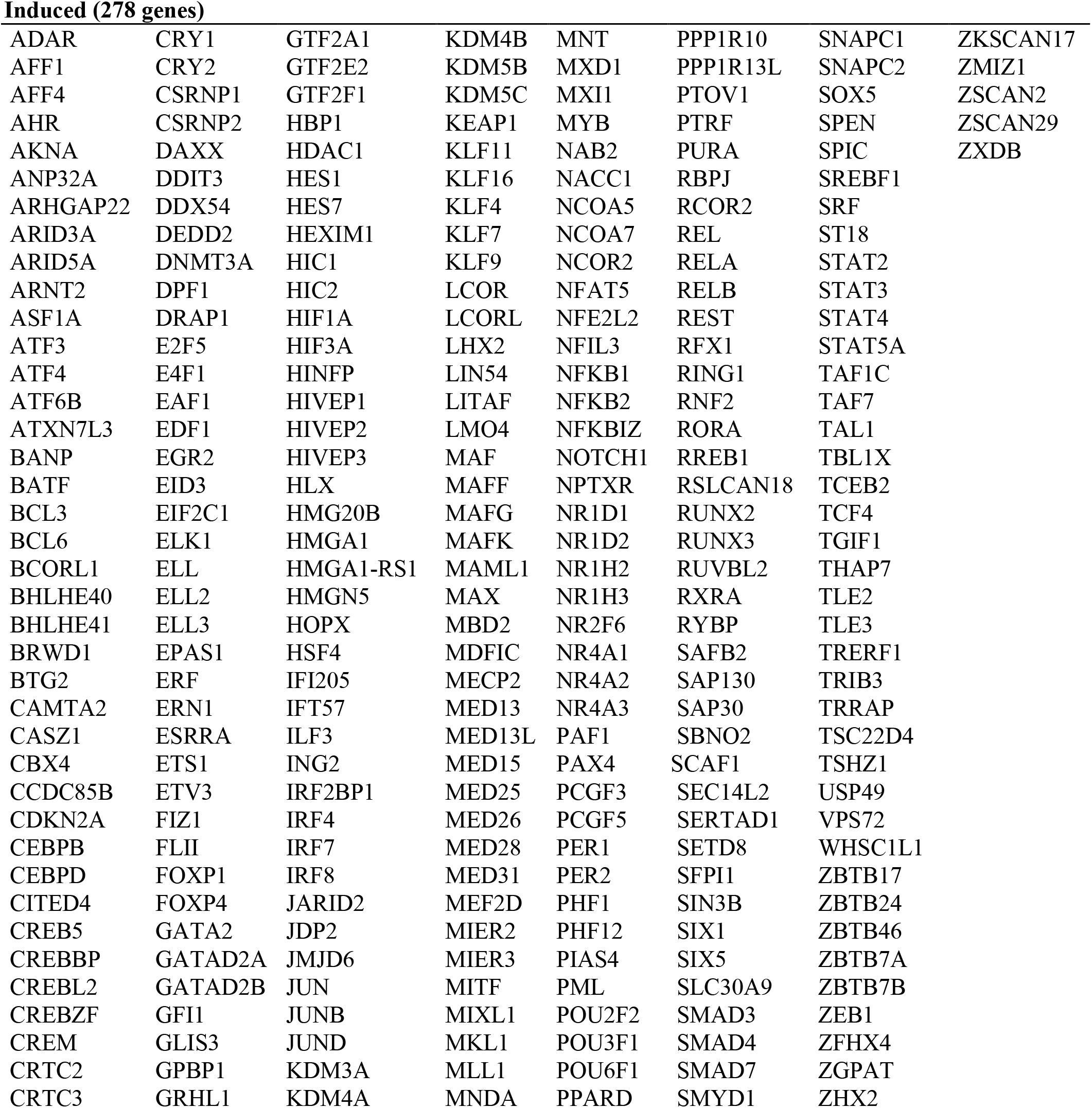
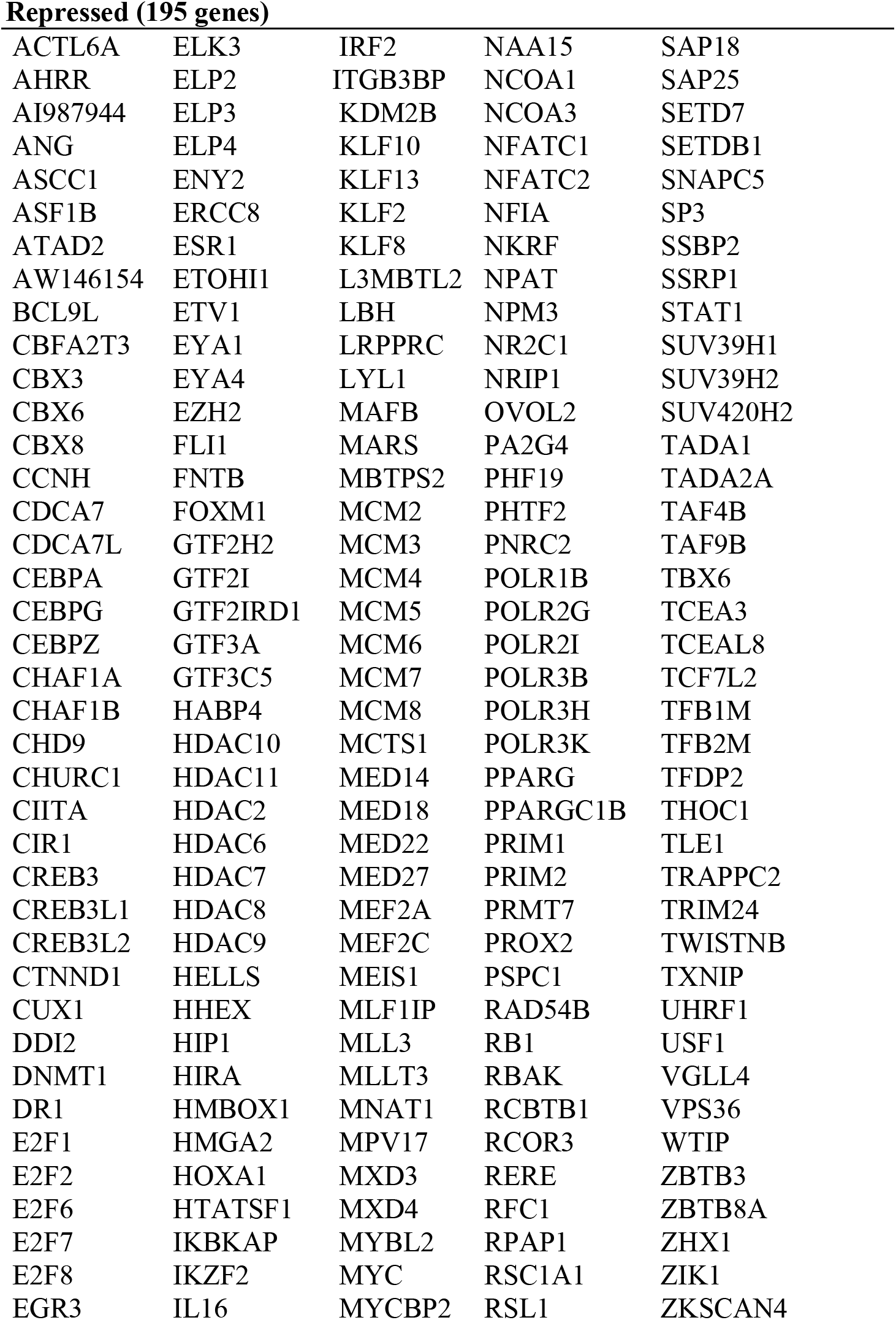
Transcriptional regulators differentially regulated in M1 activated macrophages. Genes encoding transcriptional regulators that were significantly induced or repressed by 8h M1 stimulation (p<0.05, FDR<0.05, |F.C.|>1.5) in WT bone marrow derived macrophages were identified by gene ontology analysis using the DAVID platform. Differentially regulated genes that matched the Transcription GO term in the Biological Processes GO database (accession GO:0006350) were used to generate a protein-protein interaction map using String (Supplemental figure 1). Uncharacterized zinc finger proteins (ZFPs) were omitted from analyses by String. Genes are listed below:

**Figure 2 supplemental figure 1.**
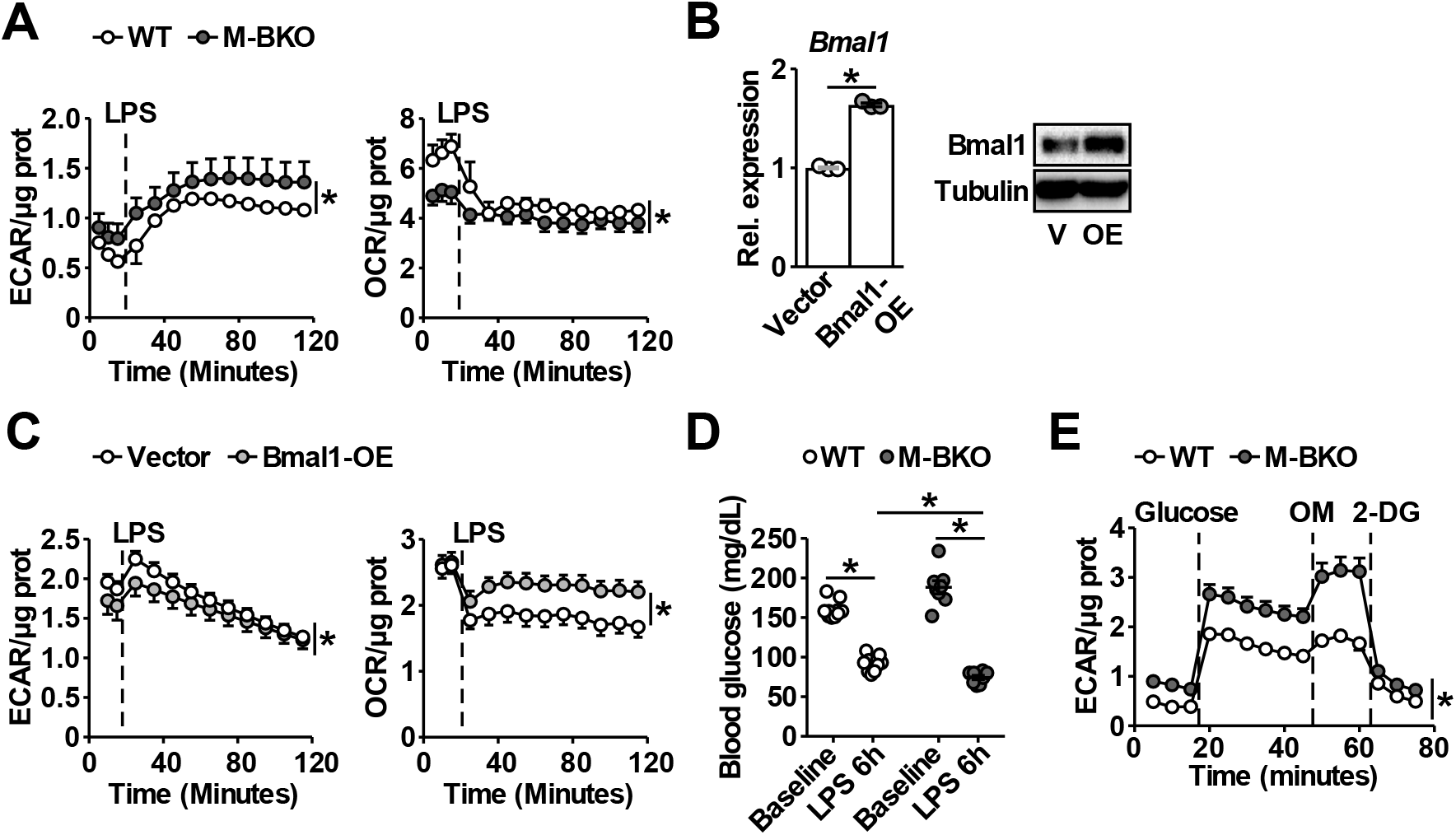
Effects of *Bmal1* gene deletion and over-expression on glycolytic versus oxidative metabolism. (**A**) Extracellular flux analysis of thioglycollate-elicited peritoneal macrophages measuring the changes in ECAR (left panel) and OCR (right panel) following LPS injection (final conc. 1 μg/mL). Assay medium contained 5 mM glucose and 1 mM pyruvate in minimal DMEM with 2% dialyzed FBS, pH 7.4. N=5 biological replicates, statistical analysis performed using 2-way ANOVA for WT vs. M-BKO across the time course. (**B**) *Bmal1* mRNA expression determined by qPCR (left) and protein levels (right) in RAW264.7 macrophage stable lines. Bmal1 OE: Bmal1 over-expressing stable line. Cells transduced with empty vector were used as the control. N=3 biological replicates for qPCR, statistical analysis performed using Student’s T test. V: vector control; OE: Bmal1 over-expression. (**C**) Measurement of the changes in ECAR (left) and OCR (right) in Ifn-γ-primed RAW264.7 stable lines after injection with LPS (final conc. 100 ng/mL). Assay medium contained 25 mM glucose and 1 mM pyruvate in minimal DMEM with 2% dialyzed FBS, pH 7.4. N=5 biological replicates, statistical analysis performed using 2-way ANOVA for control vs. Bmal1 OE across the time course. (**D**) Blood glucose levels in 4-month-old WT and M-BKO male mice before and 6 h after i.p. injection with 10 μg LPS per g body weight. N=10 mice, statistical analysis performed using Student’s T test. (**E**) Glycolytic stress test in splenic macrophages isolated from mice in (D) that were sacrificed 6 h after LPS injection. ECAR was measured before and after injection with glucose (25 mM). Maximal glycolytic rate was determined by injection of oligomycin (OM, 2 μM). Glycolysis-dependent ECAR was determined by injection with 2-deoxyglucose (2-DG, 50 mM). Assay medium contained minimal DMEM, pH 7.4. N=10 biological replicates, statistical analysis performed using 2-way ANOVA for WT vs. M-BKO across the time course. Data presented as mean ± S.E.M. *p<0.05. Experiments were repeated at least twice.

**Figure 4 supplemental figure 1.**
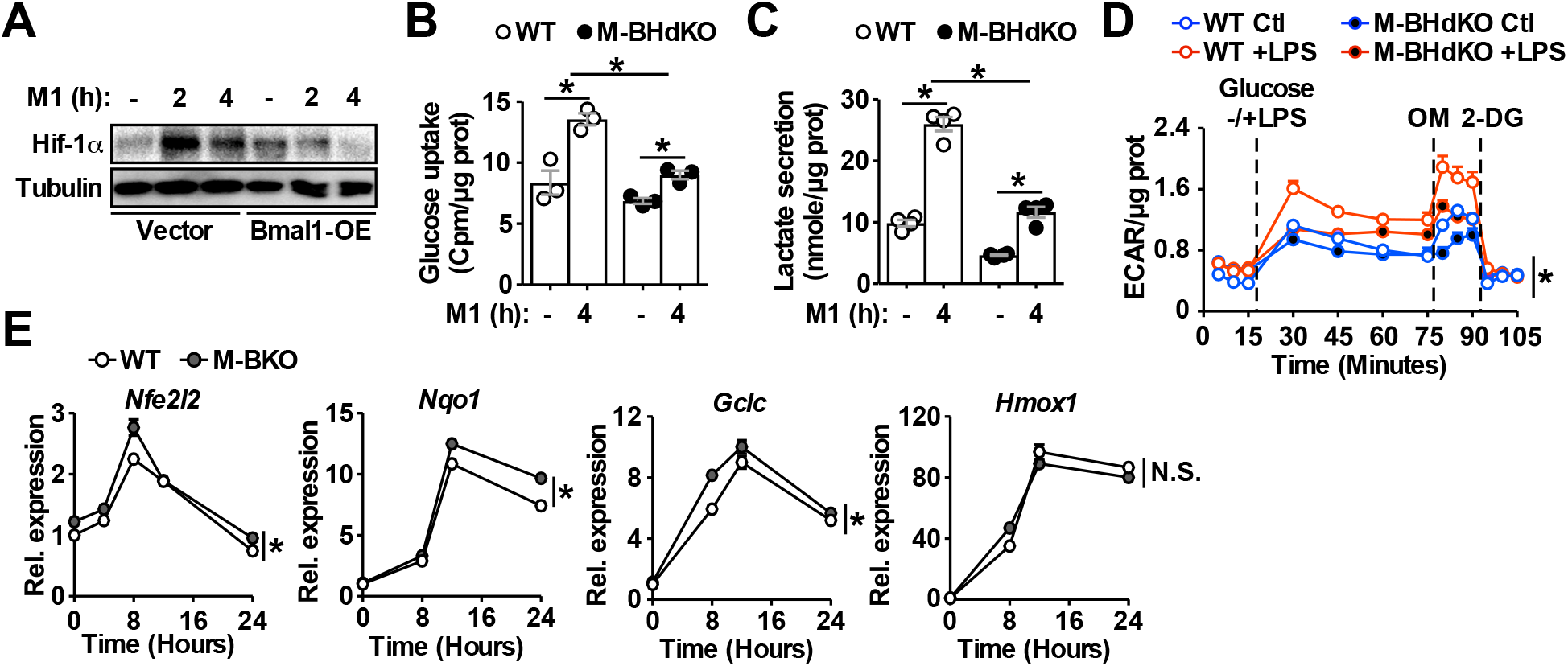
Increased oxidative stress and Hif-1α activity in M1 activated M-BKO macrophages. (**A**) Hif-1α protein levels in M1-activated (primed with 10 ng/mL Ifn-γ and stimulated with 50 ng/mL LPS) control and Bmal1 OE RAW264.7 stable lines. (**B**) and (**C**) Uptake of [^3^H]-2-deoxyglucose and lactate secretion in control or M1-activated macrophages. N=3 biological replicates, statistical analysis performed using Student’s T test. M-BHdko: myeloid-specific *Bmal1* and *Hif1a* double knockout. (**D**) Glycolytic stress test in Ifn-γ-primed macrophages measuring ECAR following glucose (25 mM) injection without or with LPS (100 ng/mL). Maximal glycolytic rate was determined by injection of oligomycin (OM, 2 μM), and glycolysis-dependent ECAR was determined by injection with 2-deoxyglucose (2-DG, 50 mM). Assay medium contained minimal DMEM with 2% dialyzed FBS, pH 7.4. N=5 biological replicates. The difference between LPS-treated WT and M-BHdKO macrophages was determined by 2-way ANOVA across the time course. (**E**) Expression of Nrf2 target genes determined by qPCR throughout a 24-hour time course following M1 stimulation. N=3 biological replicates, statistical analysis using 2-way ANOVA for WT vs. M-BKO across the time course. Data presented as mean ± S.E.M. *p<0.05. Experiments were repeated at least twice.

**Figure 5 supplemental figure 1.**
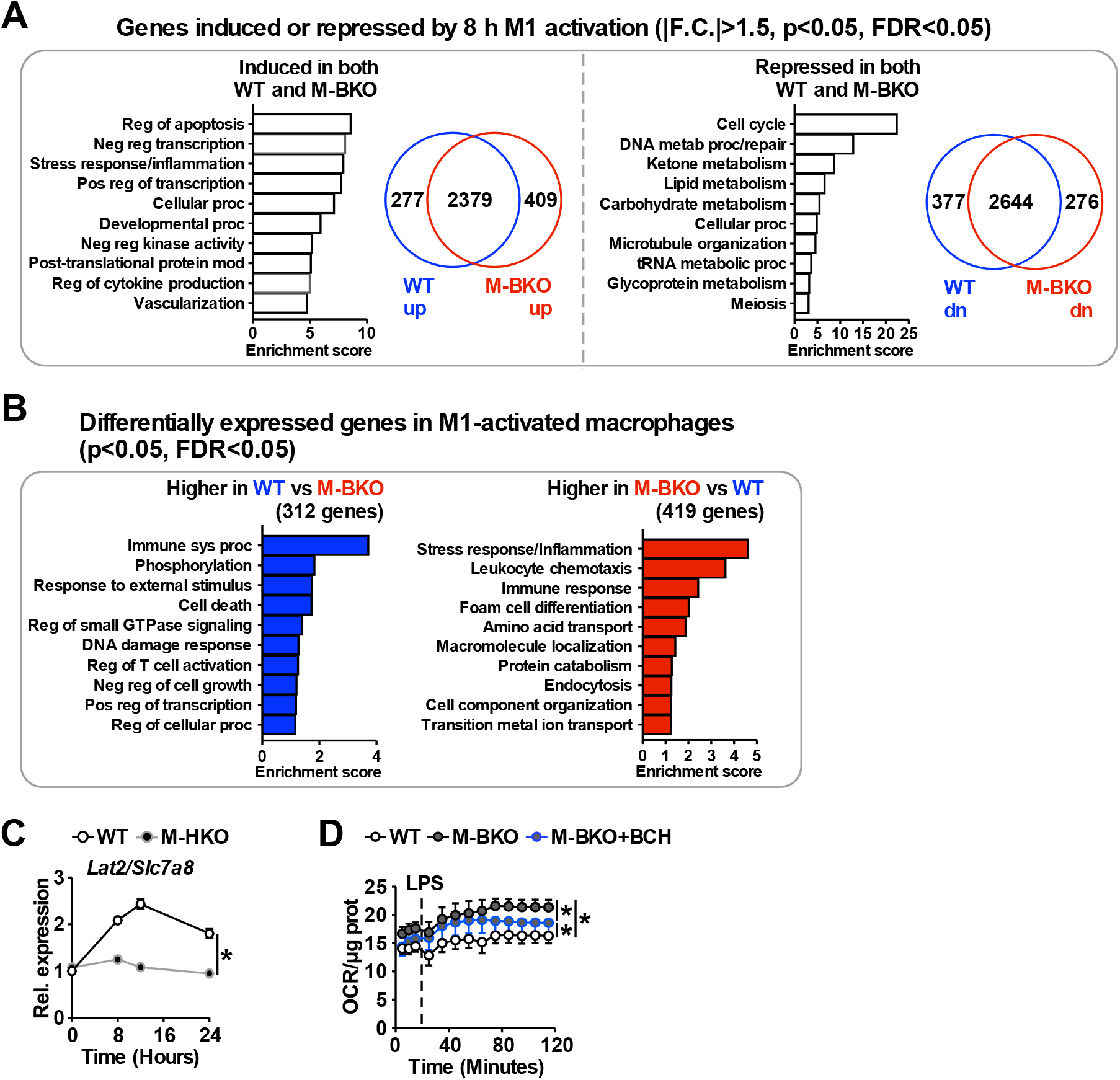
Transcriptome analysis of genes regulated by M1 stimulation in WT and M-BKO macrophages. (**A**) Functional annotation clustering by biological process and venn diagram of genes identified by RNA-seq that are mutually induced (2,379 genes, left) or suppressed (2,644 genes, right) by 8 h M1 activation in WT and M-BKO macrophages. N=3 biological replicates. F.C.: fold change. (**B**) Enriched biological processes in direct comparison of differentially expressed genes between M1-activated WT and M-BKO macrophages. (**C**) Relative expression of the neutral amino acid transporter *Slc7a8* (*Lat2*) in WT and myeloid *Hif1a* knockout (M-HKO) macrophages determined by qPCR. WT samples were from Figure 5B. N=3 biological replicates, statistical analysis performed using 2-way ANOVA for WT vs. M-HKO across the time course. (**D**) Determination of glutamine utilization by extracellular flux analysis. OCR was measured before and after LPS injection (final conc. 1 μg/mL), and assay medium contained 5 mM glutamine in minimal DMEM with 2% dialyzed FBS, pH 7.4. N=5 biological replicates, statistical analysis performed using 2-way ANOVA for WT vs. M-BKO and WT or M-BKO vs M-BKO cells co-treated with the neutral amino acid transport inhibitor 2-amino-2-norbornanecarboxylic acid (BCH, 10 mM) across the time course. Data presented as mean ± S.E.M. *p<0.05. Experiments were repeated at least twice.

**Figure 7 supplemental figure 1.**
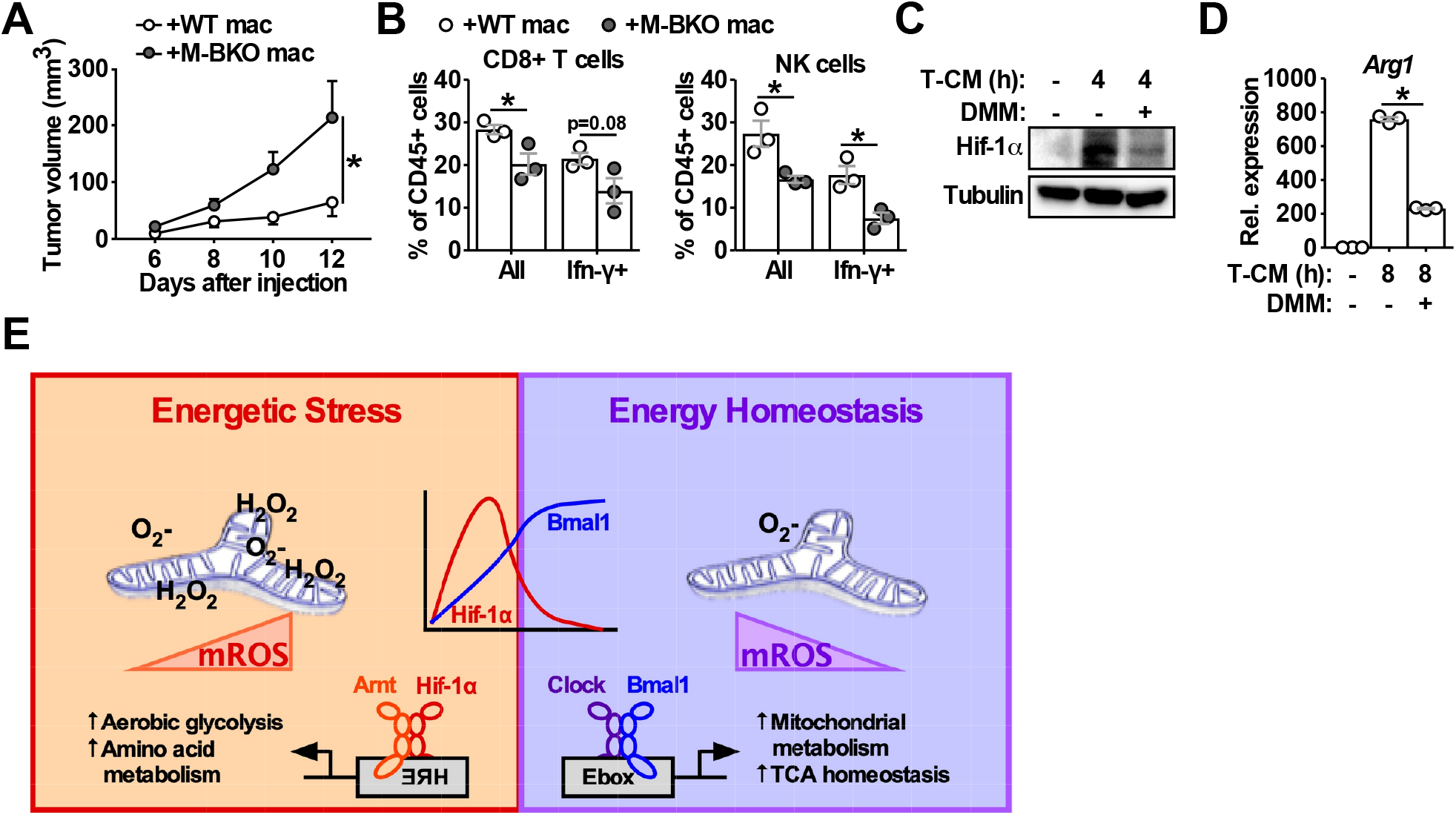
Macrophage Bmal1 regulates tumor growth in a cell-autonomous manner. (**A**) Tumor volume in M-BKO male mice co-injected with 500,000 B16-F10 cells and either 500,000 WT or M-BKO macrophages as indicated. N=6 mice, statistical analysis performed using 2-way ANOVA for WT vs. M-BKO macrophage co-injection across the time course. (**B**) Flow cytometric analyses of tumorinfiltrating CD8^+^ T cells (CD45^+^CD3^+^CD8a^+^ cells, left panel) and NK cells (CD45^+^CD3^-^NK1.1^+^ cells, right panel) stimulated *ex vivo* with phorbol 12-myristate 13-acetate and ionomycin for Ifn-γ co-staining. Tumors from (A) were pooled into three groups prior to isolation of infiltrating leukocytes for flow cytometric analyses. Statistical analysis performed using Student’s T test. Data presented as mean ± S.E.M. *p<0.05. (**C**) Hif-1α protein levels in WT macrophages treated with (from left to right) control medium or B16-F10 tumor-conditioned medium (T-CM) diluted 1:1 with control medium without or with co-treatment with 10 mM dimethyl malonate (DMM) for 4 hours. (**D**) *Arg1* gene expression in WT macrophages treated as in (C) for 8 hours. N=3 biological replicates. Statistical analysis performed using Student’s T test. Experiments were repeated at least twice. (**E**) Schematic showing the working model for Bmal1-Hif-1α crosstalk in the regulation of macrophage bioenergetics and effector functions. Both M1 and tumor-associated macrophages share similar energetically stressed states. Bmal1 preserves mitochondrial oxidative metabolism while reducing oxidative stress to modulate Hif-1α activity and support proper effector functions. Deletion of *Bmal1* in the macrophage leads to reliance on glycolytic metabolism and alternative fuel utilization, notably amino acid metabolism, which further promotes mROS production and Hif-1α protein stabilization. Increased amino acid utilization by M-BKO macrophages may lead to depletion of amino acids critical for lymphocyte activation and cytotoxic function, thereby suppressing anti-tumor immunity in the tumor microenvironment.

## REFERENCES

Andrejeva, G., & Rathmell, J. C. (2017). Similarities and Distinctions of Cancer and Immune Metabolism in Inflammation and Tumors. Cell Metab, 26(1), 49–70. doi:10.1016/j.cmet.2017.06.004

Arts, R. J., Plantinga, T. S., Tuit, S., Ulas, T., Heinhuis, B., Tesselaar, M., … Netea-Maier, R. T. (2016). Transcriptional and metabolic reprogramming induce an inflammatory phenotype in non-medullary thyroid carcinoma-induced macrophages. Oncoimmunology, 5(12), e1229725. doi:10.1080/2162402X.2016.1229725

Bell, E. L., Klimova, T. A., Eisenbart, J., Moraes, C. T., Murphy, M. P., Budinger, G. R., & Chandel, N. S. (2007). The Qo site of the mitochondrial complex III is required for the transduction of hypoxic signaling via reactive oxygen species production. J Cell Biol, 177(6), 1029–1036. doi:10.1083/jcb.200609074

Buck, M. D., Sowell, R. T., Kaech, S. M., & Pearce, E. L. (2017). Metabolic Instruction of Immunity. Cell, 169(4), 570–586. doi:10.1016/j.cell.2017.04.004

Canaple, L., Rambaud, J., Dkhissi-Benyahya, O., Rayet, B., Tan, N. S., Michalik, L., … Laudet, V. (2006). Reciprocal regulation of brain and muscle Arnt-like protein 1 and peroxisome proliferator-activated receptor alpha defines a novel positive feedback loop in the rodent liver circadian clock. Mol Endocrinol, 20(8), 1715–1727. doi:10.1210/me.2006-0052

Chouchani, E. T., Pell, V. R., Gaude, E., Aksentijevic, D., Sundier, S. Y., Robb, E. L., … Murphy, M. P. (2014). Ischaemic accumulation of succinate controls reperfusion injury through mitochondrial ROS. Nature, 515(7527), 431–435. doi:10.1038/nature13909

Christensen, H. N., Handlogten, M. E., Lam, I., Tager, H. S., & Zand, R. (1969). A Bicyclic Amino Acid to Improve Discriminations among Transport Systems. Journal of Biological Chemistry, 244(6), 1510–1520.

Colegio, O. R., Chu, N. Q., Szabo, A. L., Chu, T., Rhebergen, A. M., Jairam, V., … Medzhitov, R. (2014). Functional polarization of tumour-associated macrophages by tumour-derived lactic acid. Nature, 513(7519), 559–563. doi:10.1038/nature13490

Cramer, T., Yamanishi, Y., Clausen, B. E., Förster, I., Pawlinski, R., Mackman, N., … Johnson, R. S. (2003). HIF-1alpha is essential for myeloid cell-mediated inflammation. Cell, 112(5) 645–657.

Dai, L., Bhargava, P., Stanya, K. J., Alexander, R. K., Liou, Y.-H., Jacobi, D., … Lee, C.-H. (2017). Macrophage alternative activation confers protection against lipotoxicity-induced cell death. Molecular Metabolism, 6(10), 1186–1197. doi:10.1016/j.molmet.2017.08.001

Damiola, F., Le Minh, N., Preitner, N., Kornmann, B., Fleury-Olela, F., & Schibler, U. (2000). Restricted feeding uncouples circadian oscillators in peripheral tissues from the central pacemaker in the suprachiasmatic nucleus. Genes & development, 14(23), 2950–2961.

Doedens, A. L., Stockmann, C., Rubinstein, M. P., Liao, D., Zhang, N., DeNardo, D. G., … Johnson, R. S. (2010). Macrophage expression of hypoxia-inducible factor-1 alpha suppresses T-cell function and promotes tumor progression. Cancer Res, 70(19), 7465–7475. doi:10.1158/0008-5472.CAN-10-1439

Early, J. O., Menon, D., Wyse, C. A., Cervantes-Silva, M. P., Zaslona, Z., Carroll, R. G., … Curtis, A. M. (2018). Circadian clock protein BMAL1 regulates IL-1beta in macrophages via NRF2. Proc Natl Acad Sci U S A, 115(36), E8460–E8468. doi:10.1073/pnas.1800431115

Fukuzumi, M., Shinomiya, H., Shimizu, Y., Ohishi, K., & Utsumi, S. (1996). Endotoxin-induced enhancement of glucose influx into murine peritoneal macrophages via GLUT1. Infection and immunity, 64(1), 108–112.

Ganeshan, K., & Chawla, A. (2014). Metabolic regulation of immune responses. Annu Rev Immunol, 32, 609–634. doi:10.1146/annurev-immunol-032713-120236

Geiger, R., Rieckmann, J. C., Wolf, T., Basso, C., Feng, Y., Fuhrer, T., … Lanzavecchia, A. (2016). L-Arginine Modulates T Cell Metabolism and Enhances Survival and Anti-tumor Activity. Cell, 167(3), 829–842.e813. doi:10.1016/j.cell.2016.09.031

Hardin, P. E., & Panda, S. (2013). Circadian timekeeping and output mechanisms in animals. Current Opinion in Neurobiology, 23(5), 724–731. doi:10.1016/j.conb.2013.02.018

Hotamisligil, G. S. (2017). Foundations of Immunometabolism and Implications for Metabolic Health and Disease. Immunity, 47(3), 406–420. doi:10.1016/j.immuni.2017.08.009

Jaakkola, P., Mole, D. R., Tian, Y.-M., Wilson, M. I., Gielbert, J., Gaskell, S. J., … Ratcliffe, P. J. (2001). Targeting of HIF-α to the von Hippel-Lindau Ubiquitylation Complex by O<sub>2</sub>-Regulated Prolyl Hydroxylation. Science, 292, 468–472.

Jacobi, D., Liu, S., Burkewitz, K., Kory, N., Knudsen, Nelson H., Alexander, Ryan K., … Lee, C.-H. (2015). Hepatic Bmal1 Regulates Rhythmic Mitochondrial Dynamics and Promotes Metabolic Fitness. Cell Metabolism, 22(4), 709–720. doi:10.1016/j.cmet.2015.08.006

Kang, K., Reilly, S. M., Karabacak, V., Gangl, M. R., Fitzgerald, K., Hatano, B., & Lee, C.-H. (2008). Adipocyte-derived Th2 cytokines and myeloid PPARdelta regulate macrophage polarization and insulin sensitivity. Cell Metabolism, 7(6), 485–495. doi:10.1016/j.cmet.2008.04.002

Lamas, B., Vergnaud-Gauduchon, J., Goncalves-Mendes, N., Perche, O., Rossary, A., Vasson, M. P., & Farges, M. C. (2012). Altered functions of natural killer cells in response to L-Arginine availability. Cell Immunol, 280(2), 182–190. doi:10.1016/j.cellimm.2012.11.018

Lampropoulou, V., Sergushichev, A., Bambouskova, M., Nair, S., Vincent, Emma E., Loginicheva, E., … Artyomov, Maxim N. (2016). Itaconate Links Inhibition of Succinate Dehydrogenase with Macrophage Metabolic Remodeling and Regulation of Inflammation. Cell Metabolism, 24(1), 158–166. doi:10.1016/j.cmet.2016.06.004

Lee, C.-H., Kang, K., Mehl, I. R., Nofsinger, R., Alaynick, W. A., Chong, L.-W., … Evans, R. M. (2006). Peroxisome proliferator-activated receptor δ promotes very low-density lipoprotein-derived fatty acid catabolism in the macrophage. Proceedings of the National Academy of Sciences, 103, 2434–2439.

Li, F., Wang, Y., Zeller, K. I., Potter, J. J., Wonsey, D. R., O’Donnell, K. A., … Dang, C. V. (2005). Myc Stimulates Nuclearly Encoded Mitochondrial Genes and Mitochondrial Biogenesis. Molecular and Cellular Biology, 25(14), 6225–6234. doi:10.1128/mcb.25.14.6225-6234.2005

Liu, D., Chang, C., Lu, N., Wang, X., Lu, Q., Ren, X., … Tang, L. (2017). Comprehensive Proteomics Analysis Reveals Metabolic Reprogramming of Tumor-Associated Macrophages Stimulated by the Tumor Microenvironment. J Proteome Res, 16(1), 288–297. doi:10.1021/acs.jproteome.6b00604

Liu, S., Brown, J. D., Stanya, K. J., Homan, E., Leidl, M., Inouye, K., … Lee, C.-H. (2013). A diurnal serum lipid integrates hepatic lipogenesis and peripheral fatty acid use. Nature, 502(7472), 550–554. doi:10.1038/nature12710

Marpegán, L., Bekinschtein, T. A., Costas, M. A., & Golombek, D. A. (2005). Circadian responses to endotoxin treatment in mice. Journal of Neuroimmunology, 160(1), 102–109. doi:10.1016/j.jneuroim.2004.11.003

Masson, N., & Ratcliffe, P. J. (2014). Hypoxia signaling pathways in cancer metabolism: the importance of co-selecting interconnected physiological pathways. Cancer & metabolism, 2(1), 3–3. doi:10.1186/2049-3002-2-3

Mills, E. L., Kelly, B., Logan, A., Costa, A. S. H., Varma, M., Bryant, C. E., … O’Neill, L. A. (2016). Succinate Dehydrogenase Supports Metabolic Repurposing of Mitochondria to Drive Inflammatory Macrophages. Cell, 167(2), 457–470 e413. doi:10.1016/j.cell.2016.08.064

Mills, E. L., Ryan, D. G., Prag, H. A., Dikovskaya, D., Menon, D., Zaslona, Z., … O’Neill, L. A. (2018). Itaconate is an anti-inflammatory metabolite that activates Nrf2 via alkylation of KEAP1. Nature, 556(7699), 113–117. doi:10.1038/nature25986

Nguyen, K. D., Fentress, S. J., Qiu, Y., Yun, K., Cox, J. S., & Chawla, A. (2013). Circadian Gene Bmal1 Regulates Diurnal Oscillations of Ly6C<sup>hi</sup> Inflammatory Monocytes. Science, 341, 1483–1488.

O’Callaghan, E. K., Anderson, S. T., Moynagh, P. N., & Coogan, A. N. (2012). Long-lasting effects of sepsis on circadian rhythms in the mouse. PLoS One, 7(10), e47087. doi:10.1371/journal.pone.0047087

O’Neill, L. A., Kishton, R. J., & Rathmell, J. (2016). A guide to immunometabolism for immunologists. Nat Rev Immunol, 16(9), 553–565. doi:10.1038/nri.2016.70

Odegaard, J. I., Ricardo-Gonzalez, R. R., Goforth, M. H., Morel, C. R., Subramanian, V., Mukundan, L., … Chawla, A. (2007). Macrophage-specific PPARgamma controls alternative activation and improves insulin resistance. Nature, 447(7148), 1116–1120. doi:10.1038/nature05894

Papagiannakopoulos, T., Bauer, Matthew R., Davidson, Shawn M., Heimann, M., Subbaraj, L., Bhutkar, A., … Jacks, T. (2016). Circadian Rhythm Disruption Promotes Lung Tumorigenesis. Cell Metabolism, 24(2), 324–331. doi:10.1016/j.cmet.2016.07.001

Peek, C. B., Affinati, A. H., Ramsey, K. M., Kuo, H. Y., Yu, W., Sena, L. A., … Bass, J. (2013). Circadian clock NAD+ cycle drives mitochondrial oxidative metabolism in mice. Science, 342(6158), 1243417. doi:10.1126/science.1243417

Penny, H. L., Sieow, J. L., Adriani, G., Yeap, W. H., See Chi Ee, P., San Luis, B., … Wong, S. C. (2016). Warburg metabolism in tumor-conditioned macrophages promotes metastasis in human pancreatic ductal adenocarcinoma. Oncoimmunology, 5(8), e1191731. doi:10.1080/2162402X.2016.1191731

Robb, E. L., Hall, A. R., Prime, T. A., Eaton, S., Szibor, M., Viscomi, C., … Murphy, M. P. (2018). Control of mitochondrial superoxide production by reverse electron transport at complex I. J Biol Chem, 293(25), 9869–9879. doi:10.1074/jbc.RA118.003647

Rodriguez-Prados, J. C., Traves, P. G., Cuenca, J., Rico, D., Aragones, J., Martin-Sanz, P., … Bosca, L. (2010). Substrate fate in activated macrophages: a comparison between innate, classic, and alternative activation. J Immunol, 185(1), 605–614. doi:10.4049/jimmunol.0901698

Segawa, H., Fukasawa, Y., Miyamoto, K.-i., Takeda, E., Endou, H., & Kanai, Y. (1999). Identification and Functional Characterization of a Na+-independent Neutral Amino Acid Transporter with Broad Substrate Selectivity. Journal of Biological Chemistry, 274(28), 19745–19751.

Semenza, G. L., Roth, P. H., Fang, H. M., & Wang, G. L. (1994). Transcriptional regulation of genes encoding glycolytic enzymes by hypoxia-inducible factor 1. Journal of Biological Chemistry, 269(38), 23757–23763.

Sinclair, L. V., Rolf, J., Emslie, E., Shi, Y. B., Taylor, P. M., & Cantrell, D. A. (2013). Control of amino-acid transport by antigen receptors coordinates the metabolic reprogramming essential for T cell differentiation. Nat Immunol, 14(5), 500–508. doi:10.1038/ni.2556

Spinazzi, M., Casarin, A., Pertegato, V., Salviati, L., & Angelini, C. (2012). Assessment of mitochondrial respiratory chain enzymatic activities on tissues and cultured cells. Nat Protoc, 7(6), 1235–1246. doi:10.1038/nprot.2012.058

Steggerda, S. M., Bennett, M. K., Chen, J., Emberley, E., Huang, T., Janes, J. R., … Gross, M. I. (2017). Inhibition of arginase by CB-1158 blocks myeloid cell-mediated immune suppression in the tumor microenvironment. J Immunother Cancer, 5(1), 101. doi:10.1186/s40425-017-0308-4

Tamaru, T., Isojima, Y., van der Horst, G. T. J., Takei, K., Nagai, K., & Takamatsu, K. (2003). Nucleocytoplasmic shuttling and phosphorylation of BMAL1 are regulated by circadian clock in cultured fibroblasts. Genes to Cells, 8(12), 973–983. doi:10.1046/j.1365-2443.2003.00686.x

Tannahill, G. M., Curtis, A. M., Adamik, J., Palsson-McDermott, E. M., McGettrick, A. F., Goel, G., … O’Neill, L. A. (2013). Succinate is an inflammatory signal that induces IL-1beta through HIF-1alpha. Nature, 496(7444), 238–242. doi:10.1038/nature11986

Tsukishiro, T., Shimizu, Y., Higuchi, K., & Watanabe, A. (2000). Effect of branched-chain amino acids on the composition and cytolytic activity of liver-associated lymphocytes in rats. Journal of Gastroenterology and Hepatology, 15(8), 849–859. doi:10.1046/j.1440-1746.2000.02220.x

West, A. P., Brodsky, I. E., Rahner, C., Woo, D. K., Erdjument-Bromage, H., Tempst, P., … Ghosh, S. (2011). TLR signalling augments macrophage bactericidal activity through mitochondrial ROS. Nature, 472(7344), 476–480. doi:10.1038/nature09973

Xu, J. (2005). Preparation, Culture, and Immortalization of Mouse Embryonic Fibroblasts. Current Protocols in Molecular Biology, 70(1), 28.21.21–28.21.28. doi:doi:10.1002/0471142727.mb2801s70

Yang, X., Downes, M., Yu, R. T., Bookout, A. L., He, W., Straume, M., … Evans, R. M. (2006). Nuclear receptor expression links the circadian clock to metabolism. Cell, 126(4), 801–810. doi:10.1016/j.cell.2006.06.050

Zhang, H., Bosch-Marce, M., Shimoda, L. A., Tan, Y. S., Baek, J. H., Wesley, J. B., … Semenza, G. L. (2008). Mitochondrial Autophagy Is an HIF-1-dependent Adaptive Metabolic Response to Hypoxia. Journal of Biological Chemistry, 283(16), 10892–10903. doi:10.1074/jbc.M800102200

